# Immune determinants of leuprolide-associated cardiovascular risk in prostate cancer

**DOI:** 10.64898/2026.10.01.756122

**Authors:** Tuisha Gupta, Fanyuan Zeng, Marguerite Li, Ian Moore, Vaunita Parihar, Adithya K Yadalam, Amandeep Ahluwalia, Rendy Javier Chaparro Reyes, YuJie Chen, Jacklyn Hammons, Emily Chung, Haitong Xu, Sonia Mecorapaj, Michelle Lu, Ryan Han, Arshed A. Quyyumi, Gregory B. Lesinski, Edmund K. Waller, Brian Olson, Sagar A. Patel, Anant Mandawat, Kiranj Chaudagar

## Abstract

**Background and hypothesis:** Leuprolide, the most widely used gonadotropin-releasing hormone receptor (GnRHR) agonist for androgen deprivation therapy (ADT) in prostate cancer (PC), is associated with increased risk of cardiovascular events (CVE), but the underlying mechanisms remain poorly defined. Given the pivotal contribution of circulatory immune components to cardiovascular complications, we hypothesized that leuprolide alters the cardiac immune microenvironment and that pre-existing circulating immune phenotypes determine the patient vulnerability to CVE.

**Methods:** We integrated spatial transcriptomics of coronary plaque and cardiac tissues from leuprolide-treated mice, with longitudinal immune profiling blood leukocytes of PC patients receiving leuprolide. Spatial analyses examined treatment-associated changes in cellular composition, inferred receptor signaling, transcriptional programs and intercellular communication across coronary plaque and non-plaque cardiac compartments. Peripheral blood from 25 patients was profiled longitudinally by multiparametric flow cytometry together with measurement of circulating adipocytokines.

**Results:** Leuprolide selectively reorganized the coronary plaque microenvironment in mice, characterized by altered GnRHR- and androgen receptor-signaling in plaque fibroblasts, organization of T+B lymphoid aggregates, and enrichment of inflammatory M1-like macrophages exhibiting interferon and mesenchymal-remodeling programs. These changes accompanied by non-plaque alteration, particularly ventricular cardiomyocyte hypertrophy in murine heart. Approximately ∼44% of PC patients receiving leuprolide had CD8**^+^**T cell phenotype that associated with treatment-related CVE, including ventricular dysfunctions. A composite signature incorporating Tc1 (type 1 cytotoxic T cells), CD8**^+^**effector-memory, CD8**^+^**transitional-memory and CD8^lo^T cell populations distinguished this susceptible phenotype.

**Conclusions:** These findings establish cardiovascular immune remodeling as a framework for leuprolide-associated CVE and nominate pre-existing circulating CD8^+^T cell states as candidate biomarkers for prospective ADT-risk stratification.

**Novelty and Significance:** The mechanisms underlying cardiovascular complications of GnRH receptor agonist therapy remain incompletely understood. We combined spatial transcriptomic analysis of coronary plaque and cardiac tissue with longitudinal immune profiling of patients with prostate cancer receiving leuprolide. Leuprolide was associated with a plaque-specific immune responses, involving altered fibroblast GnRH-signaling, increased T+B lymphoid aggregates, and inflammatory M1-like macrophages with interferon and mesenchymal-remodeling programs. In contrast, histologically normal vasculature showed a predominantly M2-like macrophage response. Changes outside the plaque included adipose-stromal remodeling and an increase in hypertrophic ventricular cardiomyocytes. In patients, a pre-existing CD8**^+^**T cells had phenotype associated with treatment-related cardiovascular events, including ventricular dysfunction. A signature comprising Tc1, CD8**^+^**effector-memory, CD8**^+^**transitional-memory, and CD8**^lo^** T cell populations distinguished this immune phenotype. These observations suggest that the cardiovascular response to leuprolide is influenced by both the local tissue environment and the pre-existing systemic immune state. The circulating T-cell signature warrants prospective validation as a potential approach to identify patients who may be more susceptible to cardiovascular complications during androgen deprivation therapy.

## Introduction

Prostate cancer (PC) is the second most frequently diagnosed malignancy in men worldwide and, owing to substantial advances in screening and treatment, patients with localized disease now have an excellent long-term prognosis, with a five-year survival exceeding 99%^1^. As cancer-specific mortality continues to decline, cardiovascular disease (CVD) has emerged as a leading cause of death in this patient population, accounting for a ∼3-fold higher mortality rate than prostate cancer itself^2,3^. Consequently, minimizing treatment-associated cardiovascular toxicity or events (CVE) has become an important clinical priority. Although advanced age, socioeconomic burden, and tumor characteristics contribute to CVE risk^2,3^, these factors are largely non-modifiable at the time of cancer-treatment selection. In contrast, the choice of androgen deprivation therapy (ADT)^3^, the cornerstone of treatment for hormone-sensitive PC, represents a clinically actionable determinant of cardiovascular outcome. Therefore, defining the mechanistic contributors by which individual ADT agents differentially influence cardiovascular risk has important implications for precision oncology and cancer survivorship.

Despite producing comparable suppression of circulating testosterone and similar antitumor efficacy, gonadotropin releasing hormone (GnRH) receptor (GnRHR) agonists and antagonists have markedly different cardiovascular safety profiles^4^. In particular, leuprolide, the most commonly prescribed GnRH agonist (due to its broad clinical adoption and lower cost) has consistently been associated with ∼2-fold higher rates of major CVE than the GnRH antagonist, relugolix^4^. This discrepancy cannot be adequately explained by differences in androgen suppression, suggesting that GnRH agonists engage additional biological pathways that promote cardiovascular injury. Moreover, conventional cardiovascular risk factors, including obesity, hypertension, smoking, diabetes, dyslipidemia, and established atherosclerotic cardiovascular disease (ASCVD) risk scores that effectively predicts CVE in non-cancer patients, fail to reliably identify PC patients who preferentially develop cardiovascular complications following leuprolide therapy^5^. Together, these observations indicate that previously unrecognized mechanisms, independent of systemic androgen depletion, contribute to ADT-associated cardiovascular toxicity.

Coronary atherosclerosis is a major pathological driver of cardiovascular morbidity in patients receiving ADT^5,6^. Our recent Revolution-1 clinical trial demonstrated that leuprolide, but not relugolix, significantly increased non-calcified plaque volume (NCPV), identifying vulnerable coronary plaque as a potential substrate underlying treatment-associated CVE^5^. Because non-calcified plaques are metabolically active inflammatory lesions enriched with immune and stromal cells^7,8^, we hypothesized that GnRH agonism remodels the vascular-immune niche to promote plaque destabilization and subsequent cardiovascular injury. However, the cellular organization of these immune niches, the mechanisms coordinating immune-stromal interactions within vulnerable plaques, and whether these local tissue alterations are reflected by circulating immune biomarkers remain unknown.

Here, we integrated high-resolution spatial transcriptomics of murine coronary plaque and cardiac tissues with comprehensive immune profiling of prospectively collected blood samples from patients receiving leuprolide therapy to define the immune mechanisms underlying ADT-associated cardiovascular risk. We identify a plaque-associated immune architecture characterized by altered GnRHR- and androgen receptor-signaling in fibroblasts, organization of T+B lymphoid aggregates and accumulation of inflammatory M1-like macrophages, accompanied by cardiac alterations including ventricular cardiomyocyte hypertrophy. In patients, a distinct circulating CD8**^+^**T cell phenotype, defined with Tc1 (type 1 cytotoxic T cells), CD8**^+^**effector-memory, CD8**^+^**transitional-memory and CD8^lo^T cell populations signature was associated with treatment-associated CVE, including ventricular dysfunction. Together, these findings connect leuprolide-associated vascular immune alterations with a circulating immune phenotype and nominate pre-existing CD8**^+^**T cell states as candidate biomarkers for prospective cardiovascular risk stratification during ADT.

## Results

### Macrophages and lymphocytes define the immune architecture of coronary plaques

Given that leuprolide demonstrated a preferential increase in non-calcified plaque volume, but not calcified or low-attenuation plaque volume, in a subset of patients with PC (**Supplementary Figure 1a**)^5,7,8^, we first defined the inflammatory cellular architecture of vulnerable coronary plaques and identified populations that could contribute to treatment-associated CVE. We performed high-resolution spatial transcriptomic profiling of coronary plaque and whole heart tissue collected from Myc-CAP tumor-bearing mice treated with leuprolide during the peri-castration period and compared these tissues with those from untreated tumor-bearing controls (**Figure 1a**). To establish a permissive background of vascular inflammation and plaque formation, mice received a high-fat diet and resiquimod^9,10^, a TLR7/8 agonist, before tumor inoculation and during leuprolide treatment (**Figure 1a**). Adjacent hematoxylin and eosin (H&E)-stained heart sections were reviewed by a pathologist to delineate coronary plaque from histologically normal vasculature and surrounding cardiac compartments, enabling spatially resolved analysis using the Visium HD platform (**Figure 1b**).

**Figure 1.**
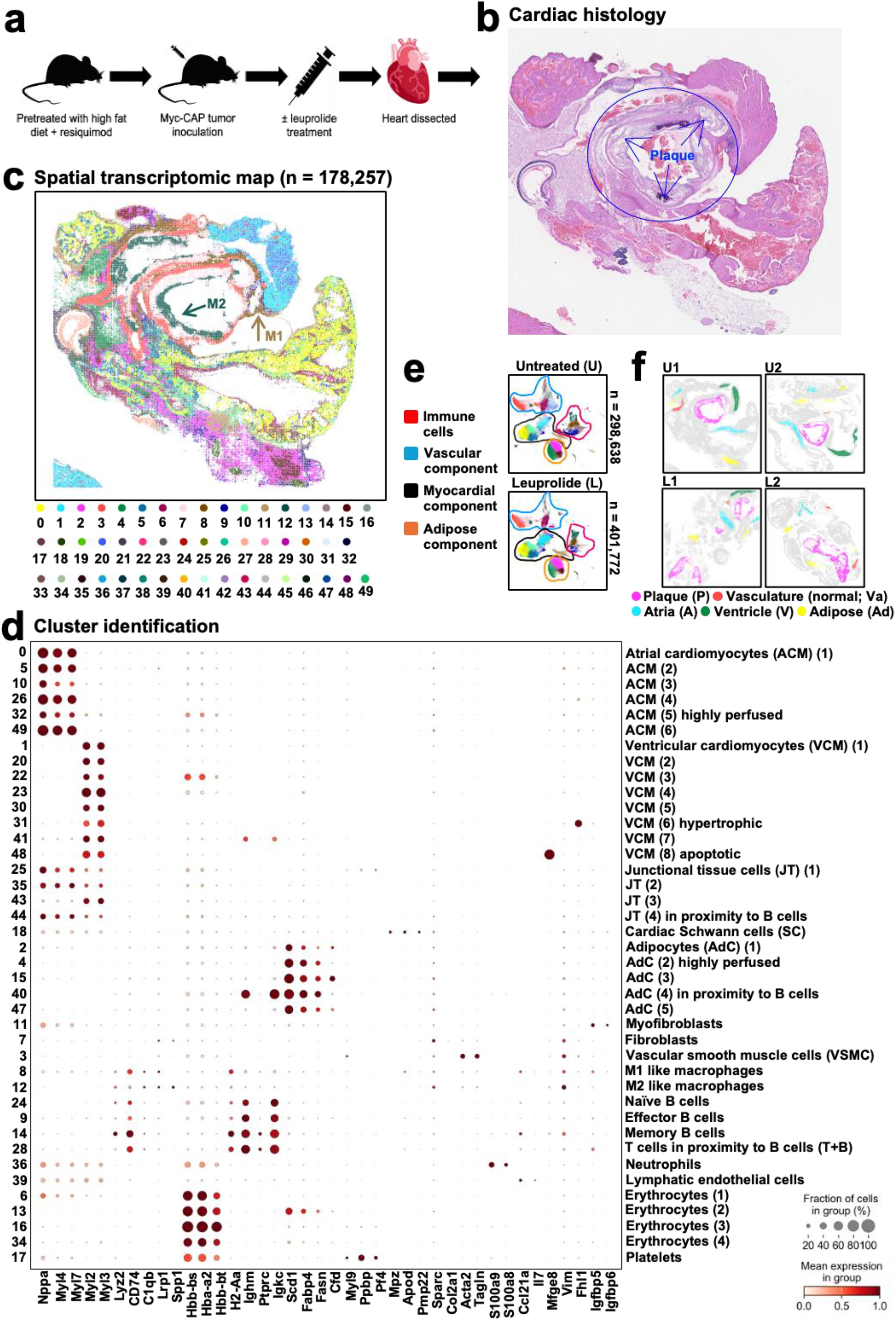
Spatial transcriptomic resolves the cellular architecture of coronary plaque and cardiac tissue. **a,** Schematic of the murine model used to investigate leuprolide-associated cardiovascular remodeling. Myc-CAP tumor-bearing mice preconditioned with a high-fat diet and resiquimod were treated with leuprolide, followed by heart collection at the peri-castration stage for spatial analysis. **b,** Representative hematoxylin and eosin (H&E)-stained section of formalin-fixed, paraffin-embedded cardiac tissue. Coronary plaque regions were identified and demarcated by pathologist. **c,** Spatial transcriptomic map generated from an adjacent cardiac section using the Visium HD platform, showing the distribution of 50 transcriptionally defined BANKSY clusters. Representing cardiac section includes 178,257 spatial profiles. Colors denote clusters and coordinates denote physical positions in the tissue section. **d,** Dot plot showing expression of representative marker genes used to annotate selected 40 transcriptionally defined BANKSY clusters retained for downstream analysis, after excluding sample-specific and technical contaminants as indicated in **Extended Figure 1e**. Dot size represents the fraction of spatial profiles expressing the indicated gene, and color intensity represents log-normalized mean expression within each population, min-max scaled independently for each gene across the displayed populations. **e,** UMAP embeddings of the 50 annotated cellular populations in untreated (U) and leuprolide-treated (L) hearts, with major compartments outlined. **f,** Pathologist-guided delineation of plaque (P), normal vasculature (Va), adipose tissue (Ad), atrial (A) and ventricular (V) regions used for subsequent compartment-resolved analyses. Data comprise 700,410 spatial profiles from four murine hearts (two untreated and two leuprolide-treated). Each spatial profile corresponds to an 8 µm Visium HD bin. U1 and U2 denote individual untreated mice; L1 and L2 denote individual leuprolide-treated mice.

Across four murine hearts, comprising two biological replicates per experimental group, spatial profiling captured 700,410 spatial bins and demonstrated consistent sample level transcriptional structure while preserving extensive cellular heterogeneity (**Supplementary Figure 1b**). Unbiased clustering identified 50 transcriptionally distinct cell populations distributed across plaque, normal vasculature and non-vascular cardiac compartments in both treatment and control conditions (**Figure 1c and Supplementary Figure 1c**). Within pathologist-defined plaque regions, we identified 13 major cellular clusters representing vascular smooth muscle cells (VSMC), fibroblasts, macrophages, B cells, T cell-enriched lymphoid populations, erythrocytes and platelets (**Figure 1c, d**). VSMC were characterized by expression of *Acta2*, *Tagln* and *Vim*^11,12^ and demarcated both plaque-containing and histologically normal vascular structures from surrounding myocardium. Fibroblasts, positioned adjacent to VSMC and preferentially toward the luminal aspect of the vascular wall, were defined by expression of *Sparc* and *Vim*^11,12^ (**Figure 1c, d**). Macrophages constituted a prominent component of the plaque immune microenvironment and were identified by expression of *C1qb*, *Lyz2* and *Lrp1*^11,12^. Based on transcriptional programs associated with inflammatory versus (*vs*) reparative states, macrophages were further classified as M1-like populations enriched for *H2-Aa* and *Cd74* or M2-like populations characterized by *Spp1* and *Vim*^11–14^ (**Figure 1c, d**). Both macrophage states were highly represented within plaque regions and formed dense spatial accumulations relative to non-plaque vascular compartments, consistent with a macrophage-rich inflammatory niche.

Multiple B cell populations were also detected within plaque and broadly characterized by expression of *Ighm* and *Igkc*^15,16^. Differential expression of *Cd74*, *H2-Aa* and *Ccl21*^15,16^ further resolved these populations into naïve-, memory- and effector-like B cell states (**Figure 1c, d**). T cell-enriched regions were identified by *Cd8a*^15^ and *Themis*^17^ expression. Notably, these regions showed substantial spatial overlap with transcriptional signatures associated with memory B cells, limiting confident separation of the two populations at the local spatial resolution. We therefore designated these mixed lymphoid regions as T+B clusters or lymphoid aggregates, reflecting their close spatial organization rather than a discrete hybrid cellular phenotype (**Figure 1c, d and Supplementary Figure 1d**). Platelets, defined by *Myl9*, *Ppbp* and *Pf4*^11,12^, and four erythrocyte populations expressing *Hba-A2*, *Hbb-bs* and *Hbb-bt*^11,12^ were readily detected but were considered predominantly intravascular components and were excluded from subsequent plaque-interaction analyses. Outside plaque regions, we additionally identified neutrophils expressing *S100a8* and *S100a9*^11,12^, as well as a small population of lymphatic endothelial-like cells expressing *Ccl21a* and *Il7*^11,12^ (**Figure 1c, d**).

We next examined the major non-vascular cardiac populations to establish whether leuprolide-associated changes extended beyond coronary plaque. Cardiomyocytes formed 14 transcriptionally distinct clusters comprising six atrial and eight ventricular populations. Atrial cardiomyocytes were characterized by *Nppa*, *Myl4* and *Myl7*^18^, whereas ventricular cardiomyocytes expressed *Myl2* and *Myl3*^18^ (**Figure 1c, d**). Cardiomyocytes with hypertrophic or apoptotic transcriptional features were further distinguished by enrichment of *Fhl1* and *Mfge8*^19^, respectively. Four junctional tissue populations co-expressed atrial and ventricular cardiomyocyte programs. We also identified myocardial myofibroblasts expressing *Igfbp5* and *Igfbp6*^18^, which were transcriptionally distinct from vascular fibroblasts, and cardiac Schwann-like cells, characterized by *Mpz*, *Apod* and *Pmp22*^20^.

Given the established contribution of cardiac adipose tissue to cardiovascular inflammation, we further resolved five adipocyte populations defined by *Scd1*, *Fabp4*, *Fasn* and *Cfd*^21^ (**Figure 1c, d**). One adipocyte cluster showed substantial overlap with naïve B cell signatures, consistent with close spatial proximity between adipocytes and lymphoid cells. Two additional clusters restricted to a single heart were excluded from downstream analyses as sample-specific populations, and small gastrointestinal and respiratory tissue signatures^22,23^ introduced during tissue collection were similarly treated as technical contaminants (**Supplementary Figure 1e**). Together, these spatial analyses establish a comprehensive cellular map of the murine heart and reveal coronary plaque as a distinct immune-rich niche characterized by dense macrophage populations and spatially organized lymphoid aggregates. This framework provided the basis for determining how leuprolide selectively remodels immune interactions within vulnerable plaque.

### Leuprolide selectively enriches inflammatory macrophages and lymphoid aggregates within coronary plaque

Whole heart analysis revealed no significant differences in the overall abundance of major cellular populations between leuprolide-treated and untreated mice (**Figure 1e and Supplementary Figure 2a**), potentially reflecting the cellular heterogeneity and anatomical variation across cardiac sections. We therefore performed compartment-resolved analyses based on five regions independently delineated by a pathologist on adjacent histological sections: coronary plaque (P), normal vasculature (Va), adipose tissue (Ad), atria (A) and ventricle (V) (**Figure 1f**). This spatially restricted approach enabled us to distinguish treatment-associated alterations within vascular lesions from changes occurring across the surrounding cardiac tissue.

Within coronary plaques, leuprolide treatment increased the abundance of inflammatory M1-like macrophages compared with untreated controls, whereas no corresponding enrichment was observed in histologically normal vasculature (**Figure 2a and Supplementary Figure 2b**). M2-like macrophages were increased in both plaque and normal vasculature following leuprolide exposure, consistent with a concomitant reparative macrophage response to vascular inflammation. Notably, T+B lymphoid aggregates (T+B clusters) were selectively increased within leuprolide-exposed plaques but remained unchanged in normal vasculature, accompanied by a trend toward increased memory B cells (**Figure 2a and Supplementary Figure 2b**). Conversely, effector B cells were reduced in both plaque and normal vasculature. Other immune populations, fibroblasts and VSMC showed no substantial treatment-associated differences in abundance within these compartments. Plaque and normal vascular regions were also devoid of appreciable cardiomyocyte or adipocyte signatures (**Supplementary Figure 2c, d**), supporting the anatomical specificity of these vascular immune alterations.

**Figure 2.**
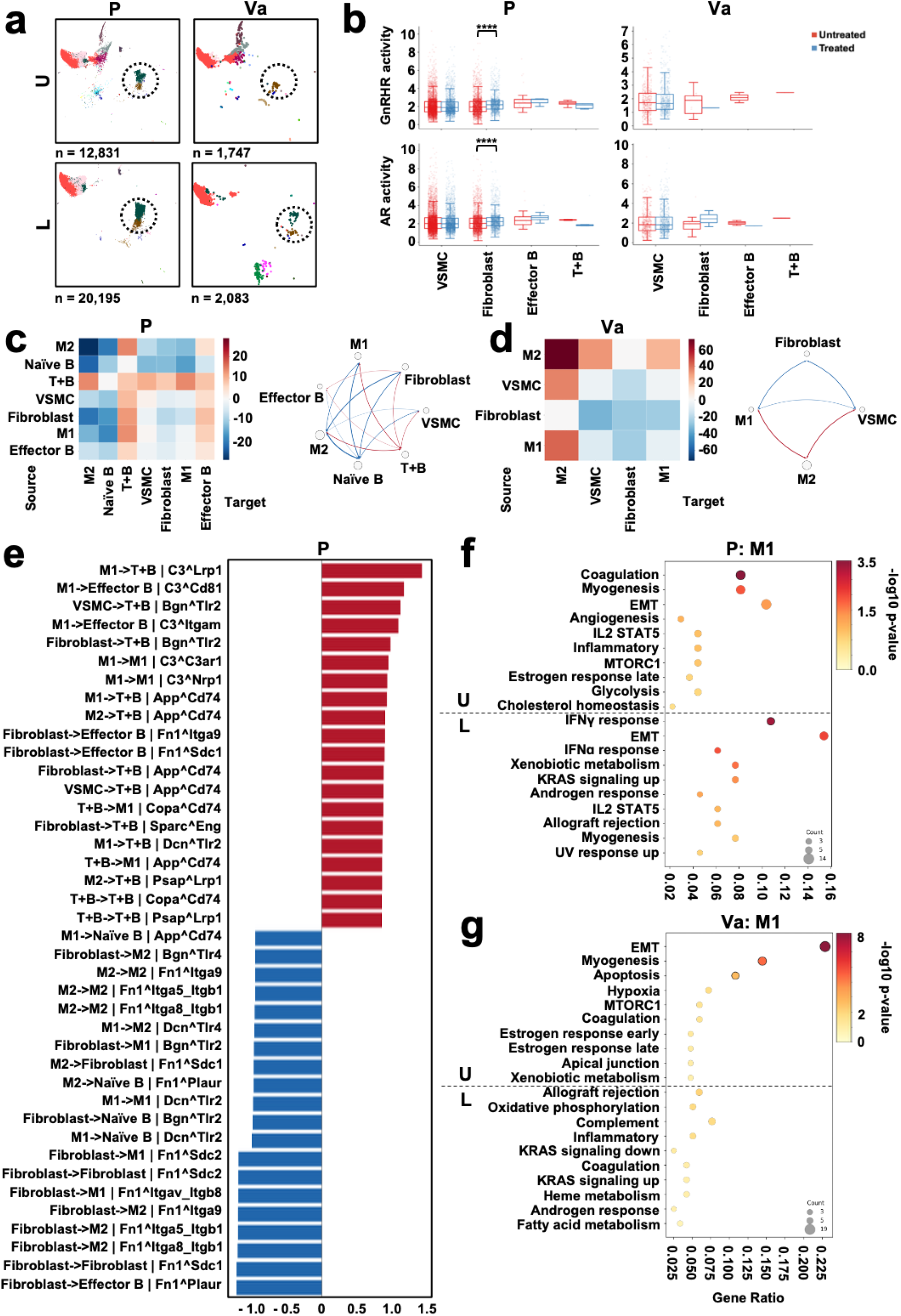
Leuprolide orchestrates fibroblast-lymphoid-macrophage axis within coronary plaque. **a,** UMAP embeddings of transcriptionally defined immune and stromal populations within pathologist-delineated coronary plaque (P) and histologically normal vasculature (Va) from untreated (U) and leuprolide-treated (L) hearts. Experimental design is shown in **Figure 1a**. **b,** Univariate linear model scores androgen receptor (AR)- and gonadotropin-releasing hormone receptor (GnRHR)-pathways transcription program across cell populations within plaque and normal vasculature. Points denote individual spatial bin with a score greater than 0. Boxes show the median and interquartile range (IQR), and whiskers extend to 1.5 × IQR. Distributions were compared within each cluster between untreated and treated arms using two-sided Mann-Whitney U tests, with Benjamini-Hochberg (BH) correction within each region. ****p<0.0001. **c,d,** Differential LIANA-predicted cell-cell communication networks in leuprolide-treated tissues compared to untreated within plaque (**c**) and normal vasculature (**d**) regions, displayed as heatmaps (left) and circos plots (right). Heatmaps show the summed difference in LIANA consensus magnitude for each sender-receiver pair, circos plots show the sender-receiver edges with the largest absolute differences. Arrows point from sender to receiver, edge width is proportional to the absolute difference, and node size represents total incident absolute difference. Red denotes increased and blue denotes decreased predicted communication following leuprolide treatment relative to untreated controls. **e,** Top differential ligand-receptor interactions within coronary plaque following leuprolide treatment. Bars show treated-minus-untreated LIANA consensus magnitude for the top 20 treated-high (red color) and top 20 untreated-high (blue color) sender-receiver ligand-receptor combinations. **f,g,** Hallmark pathway enrichment of M1-like macrophages (M1) within coronary plaque (**f**) and normal vasculature (**g**) from untreated and leuprolide-treated mice hearts using Over-Representation Analysis (ORA). Horizontal position denotes gene ratio (overlapping genes divided by query-set size), point size denotes overlap count and colour denotes -log10(raw ORA p-value). Black outline denotes BH-adjusted ORA p< 0.05. Analyses included 33,026 and 3,830 spatial profiles within pathologist-defined plaque and normal vascular regions, respectively, obtained from four murine hearts (two untreated and two leuprolide-treated). Quantitative data are presented as mean±s.e.m.

We next investigated whether leuprolide was associated with altered GnRHR-signaling within specific plaque-resident populations. Compared with endogenous GnRH-associated inferred activity in untreated mice, leuprolide markedly increased the (inferred) GnRHR-signaling activity^24^ in plaque fibroblasts (**Figure 2b**). In contrast, fibroblasts within normal vasculature exhibited reduced GnRHR-signaling activity following treatment, revealing a striking compartment-dependent response to GnRHR agonism. Because leuprolide simultaneously suppresses systemic androgen signaling, we inferred androgen receptor (AR)-signaling activity^25^ across the same populations. Unexpectedly, the inferred AR-signaling activity was increased in plaque fibroblasts following leuprolide treatment, whereas no significant alterations were detected among major cellular populations within normal vasculature (**Figure 2b**). Thus, leuprolide exposure was associated with coordinated GnRHR and AR transcriptional activation specifically within plaque-resident fibroblasts.

We therefore examined whether this fibroblast state was accompanied by remodeling of intercellular communication within plaque. In leuprolide-exposed plaques, T+B lymphoid aggregates emerged as a prominent communication hub, receiving computationally predicted signals from multiple plaque-resident populations, including fibroblasts, while transmitting signals to surrounding cells, including M1-like macrophages (**Figure 2c**). Effector B cells primarily exhibited a signal-receiving phenotype, although their predicted communication was substantially weaker than that of the T+B aggregates. By contrast, normal vasculature displayed a distinct communication architecture dominated by M2-like macrophages, which both received and transmitted signals across multiple vascular populations following leuprolide exposure (**Figure 2d**). Collectively, these findings identify a spatially restricted plaque response to leuprolide characterized by altered fibroblast signaling, enrichment of T+B lymphoid aggregates and accumulation of inflammatory M1-like macrophages, contrasting with a predominantly M2-like macrophage response in normal vasculature.

### Lymphoid aggregates organize inflammatory programs associated with plaque destabilization

To define the fibroblast programs underlying leuprolide-induced remodeling of the coronary plaque immune niche, we integrated ligand-receptor inference, pathway enrichment and transcription factor activity analyses. Ligand-receptor analysis revealed substantial reorganization of fibroblast-mediated communication following leuprolide exposure. *Fn1* and *Bgn*, extracellular matrix-associated genes implicated in plaque structural integrity^26^, were expressed by plaque fibroblasts and showed broadly reduced predicted ligand-receptor interactions, including *Itgb1*, *Itgb8*, *Itga9*, *Sdc1*, *Sdc2*, *Tlr2* and *Tlr4*, on macrophage and naïve B-cell populations in leuprolide-treated mice compared with controls (**Figure 2e**). In contrast, selected interactions were retained or enhanced within lymphoid populations, including *Bgn-Tlr2* signaling involving T+B aggregates and *Fn1-Itga9/Sdc1* interactions involving effector B cells (**Figure 2e**). These changes coincided with increased predicted interactions involving *C3* and *Cd74*^27^, molecular programs associated with complement activation and inflammatory antigen presentation, respectively. Normal vasculature displayed a distinct response characterized by attenuation of fibroblast-derived *Fn1* and *Bgn* interactions and increased *C3*-associated signaling but lack of *Cd74*-associated program (**Supplementary Figure 3a**), indicating that leuprolide differentially reorganizes extracellular matrix and immune communication, partly due to lymphoid-associated program within plaque compared to normal vasculature.

We next examined pathway level changes accompanying this altered cellular communication. At baseline, plaque fibroblasts exhibited greater epithelial/endothelial-to-mesenchymal transition (EMT), complement, coagulation and apoptosis gene expression signatures than corresponding normal vasculature, consistent with the inflammatory environment established by high-fat diet and resiquimod exposure^9,10^ (**Supplementary Figure 3b**). Leuprolide reduced several of these programs in both vascular compartments; however, EMT-associated transcriptional activity persisted within plaque while declining in normal vasculature (**Supplementary Figure 3b**), suggesting selective maintenance of a fibroblast remodeling phenotype within the plaque microenvironment. Further, T+B lymphoid aggregates showed an increased apical junction-associated transcriptional programs following leuprolide treatment, including integrins and cellular homing receptors that may facilitate their spatial organization and interactions with neighboring plaque populations (**Supplementary Figure 3c**). Consistent with persistent plaque remodeling, Sox17^28,29^-associated regulatory networks linked to mesenchymal programs were enriched in plaque fibroblasts, whereas the immune-regulatory transcription factors Irf1 and Ciita were preferentially activated within T+B aggregates following leuprolide exposure (**Supplementary Figure 3d**). These lymphoid alterations were accompanied by a distinct inflammatory state in plaque macrophages. M1-like macrophages from leuprolide-exposed plaques exhibited increased IFNα- and IFNγ-response programs together with enhanced EMT-associated transcriptional activity (**Figure 2f**). These inflammatory programs were not observed in M1-like macrophages within normal vasculature (**Figure 2g**). Conversely, EMT-associated activity in plaque M2-like macrophages was prominent in untreated mice but diminished following leuprolide treatment (**Supplementary Figure 4a**). Within normal vasculature, M2-like macrophages exhibited minimal IFNα-response activity after leuprolide exposure (**Supplementary Figure 4b**), consistent with a predominantly reparative response. Together, these analyses identify a plaque-selective program characterized by T+B lymphoid aggregates, inflammatory M1-like macrophages and altered fibroblast communication, coupling changes in extracellular matrix-associated interactions with interferon and persistent mesenchymal programs associated with plaque destabilization.

### Leuprolide alters an adipocyte-B cell stromal remodeling program outside coronary plaque

Given the well-established contribution of adipose tissue-B cell interactions to vascular inflammation^30^, we next hypothesized that leuprolide remodels cardiac immune-stromal niche beyond the coronary plaque that potentially contributes to adverse cardiac consequences. Compartment-specific analysis revealed alterations in M1-like macrophages and B cell populations within cardiac adipose tissues of leuprolide-treated mice compared with untreated controls (**Figure 3a and Supplementary Figure 2b**). Memory and effector B cell populations were enriched in this compartment, whereas naïve B cells were reduced following leuprolide exposure, consistent with their reduction within vascular regions (**Supplementary Figure 2b**). Notably, neither T+B lymphoid aggregates nor M2-like macrophages showed substantial changes in abundance outside plaque (**Supplementary Figure 2b**), supporting spatial restriction of the fibroblast-lymphoid aggregate-M1-like macrophage program identified within the coronary plaque niche.

**Figure 3.**
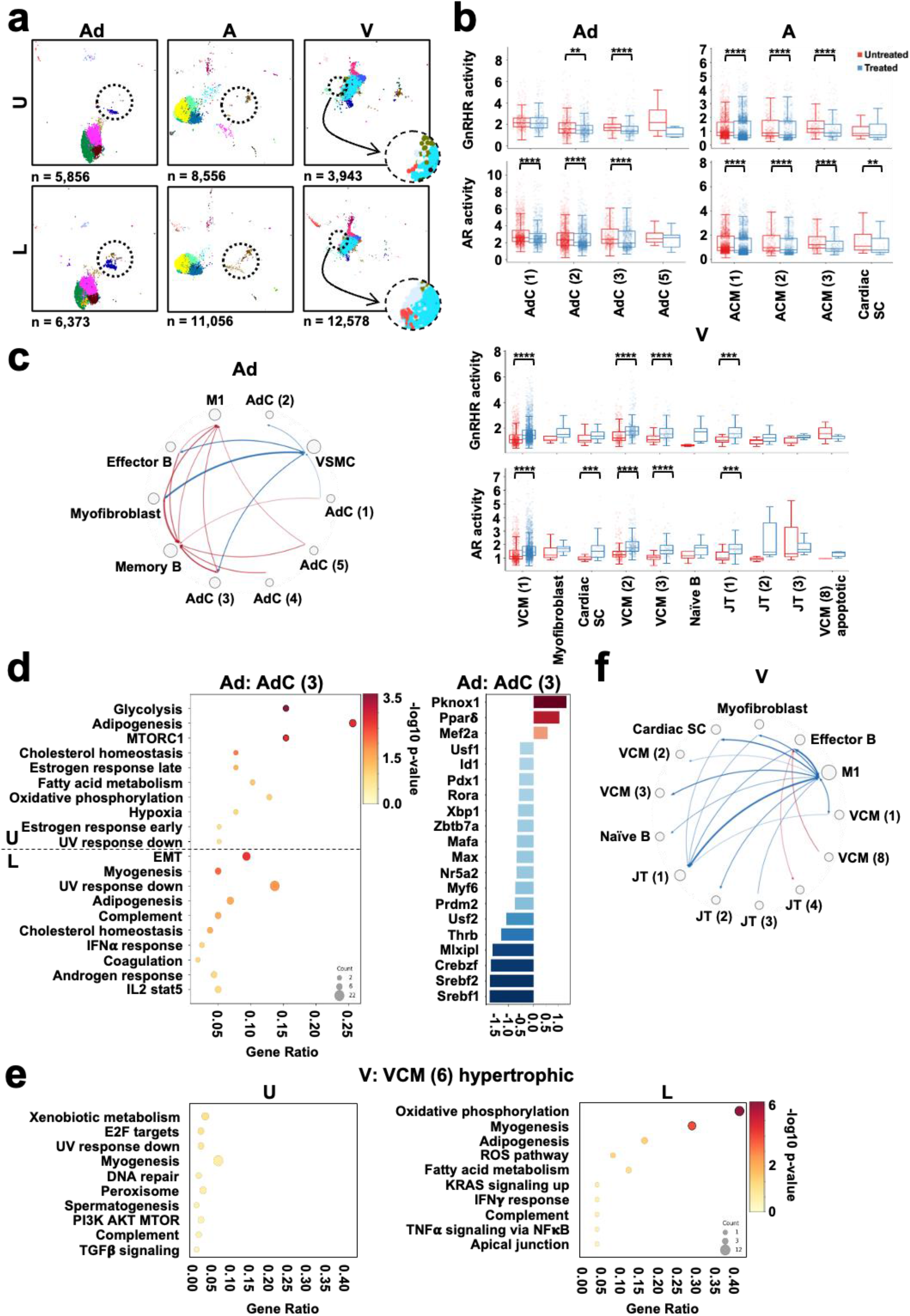
Leuprolide induces adipose-stromal remodeling and ventricular cardiomyocyte hypertrophy. **a,** UMAP embeddings of transcriptionally defined populations within pathologist-delineated cardiac adipose tissue (Ad), atrial (A) and ventricular (V) compartments from untreated (U) and leuprolide-treated (L) hearts. Experimental design is shown in **Figure 1a**. **b,** Univariate linear model (ULM) scores androgen receptor (AR)-and gonadotropin-releasing hormone receptor (GnRHR)-pathways transcription program across adipocyte (Adc), atrial cardiomyocyte (ACM), ventricular cardiomyocyte (VCM) and junctional tissue (JT) populations in untreated and leuprolide-treated hearts. Points denote individual spatial bin with a score greater than 0. Boxes show the median and interquartile range (IQR), and whiskers extend to 1.5 × IQR. Distributions were compared within each cluster between untreated and treated arms using two-sided Mann-Whitney U tests, with Benjamini-Hochberg (BH) correction within each region. **p<0.01 and ****p<0.0001. **c,** Differential LIANA-predicted cell-cell communication within cardiac adipose tissue following leuprolide treatment, displayed as a circos plot, which shows the sender-receiver edges with the largest absolute differences. Arrows point from sender to receiver, edge width is proportional to the absolute difference, and node size represents total incident absolute difference. Red denotes increased and blue denotes decreased predicted communication in leuprolide-treated relative to untreated tissue; arrows indicate the direction of inferred signaling from sender to receiver populations. **d,** Hallmark pathway enrichment analysis and CollecTRI/ULM transcription factor (TF) activity differences of adipocyte subset 3 (Adc (3)). TF bars show the signed difference in mean profile-level ULM activity (treated minus untreated) for the 20 TF with the largest absolute differences. **e,** Hallmark pathway enrichment analysis of ventricular cardiomyocyte subset 6 (VCM (6)), transcriptionally characterized by a hypertrophic phenotype, in untreated and leuprolide-treated hearts. **f,** Differential predicted cell-cell communication among ventricular cellular populations following leuprolide treatment, displayed as a circos plot. Red and blue indicate increased and decreased predicted communication, respectively, in leuprolide-treated relative to untreated mice ventricles. Analyses included 12,229, 19,612 and 16,521 spatial profiles within adipose, atrial and ventricular regions, respectively, obtained from four murine hearts (two untreated and two leuprolide-treated). Quantitative data are presented as mean±s.e.m.

Beyond these immune alterations, adipocyte subsets 3 and 4 were markedly enriched following leuprolide treatment, with adipocyte subset 4 localized in close proximity to B cells (**Supplementary Figure 2d**). To minimize potential confounding from overlapping adipocyte and B cell transcriptional signals, we therefore examined the transcriptional state and cellular interactions of adipocyte subset 3. Following leuprolide treatment, both inferred AR and GnRHR-signaling activities were reduced in adipocyte subset 3, as well as in additional adipocyte populations (**Figure 3b**). Concomitantly, predicted communication between adipocyte subset 3 and memory B cells increased in treated mice, involving *C3*-associated signaling and *Lpl-Lrp1* interactions (**Figure 3c and Supplementary Figure 5a**). Pathway and transcription factor activity analyses further revealed a treatment-associated shift in adipocyte and B cell states. At baseline, adipocyte subset 3 exhibited prominent adipogenesis accompanied by glycolytic and mTORC1-associated programs (**Figure 3d**), whereas memory B cells displayed transcriptional enrichment for fatty acid metabolism (**Supplementary Figure 5b**). Following leuprolide exposure, both populations exhibited increased EMT-associated transcriptional programs, with increased EMT-associated activity and transcription factor networks involving Pknox1, Smad2, Smad7 and Snai1^31–33^ (**Figure 3d and Supplementary Figure 5b**). These changes indicate a coordinated shift from metabolic toward EMT-associated transcriptional programs within the adipocyte-B cell niche.

Furthermore, leuprolide increased predicted communication between effector B cells and myofibroblasts (**Figure 3c**). Although myofibroblast abundance was not substantially altered within adipose tissue, these cells markedly accumulated within the atrial compartment (**Figure 3a and Supplementary Figure 2c**). Atrial cardiomyocytes concurrently exhibited reduced the inferred AR and GnRHR-signaling activities following treatment (**Figure 3b**). Myofibroblasts, which displayed adipogenesis and fatty acid metabolism programs at baseline, lost these metabolic features following leuprolide exposure and acquired an increased Pknox1-associated regulatory activity (**Supplementary Figure 5b**). This transition was accompanied by enhanced predicted communication between myofibroblasts and M1-like macrophages within cardiac adipose tissue (**Figure 3c**). Collectively, these findings identify a spatially distinct adipocyte-B cell-myofibroblast remodeling program associated with coordinated attenuation of AR and GnRHR signaling following leuprolide exposure. This non-plaque response contrasts with the GnRHR agonism-driven inflammatory lymphoid-macrophage program within coronary plaque and suggests that leuprolide simultaneously remodels multiple cardiac tissue niches that may contribute to cardiovascular toxicity through distinct mechanisms.

### Leuprolide is associated with ventricular cardiomyocyte hypertrophy

Given the reported effects of GnRHR-signaling on cardiomyocyte contractility^34^, we next examined whether leuprolide directly altered cardiomyocyte states. Cardiomyocyte profiling revealed an approximately 4-fold increase in hypertrophic ventricular cardiomyocytes (subset 6) compared with untreated controls (**Figure 3a and Supplementary Figure 2c**). These cells exhibited increased oxidative phosphorylation and myogenesis programs, consistent with a hypertrophic transcriptional state (**Figure 3e**). Although the inferred AR and GnRHR-signaling increased across several ventricular populations, hypertrophic cardiomyocytes lacked detectable activity of either pathway (**Figure 3b**). Moreover, they showed limited predicted communication with other cardiomyocyte populations (**Figure 3f**). Thus, leuprolide-associated ventricular hypertrophy occurs without detectable cell-intrinsic AR or GnRHR activation, suggesting that cardiac remodeling may arise secondary^35,36^ to treatment-induced alterations in the surrounding vascular or cardiac microenvironment.

### A subset of patients with primary PC exhibits a circulating CD8+T cell-enriched phenotype

Having identified distinct immune and stromal programs associated with leuprolide exposure in coronary plaque and non-plaque cardiac compartments in a murine model of PC, we next asked whether these tissue-level alterations could identify the patients having CVE risk following leuprolide. We profiled peripheral blood from 25 patients with primary PC who received leuprolide therapy (**Figure 4a**). Prospectively collected blood samples spanning baseline and on-treatment time points were analyzed using multiparametric flow cytometry incorporating 29 markers to resolve 26 immune populations, together with measurement of 15 circulating adipocytokines. Clinical follow-up captured CVE, including cardiomyopathy, electrocardiographic abnormalities and angina (**Supplementary Table 1**), enabling subsequent evaluation of relationships between circulating immune phenotypes and cardiovascular outcomes.

**Figure 4.**
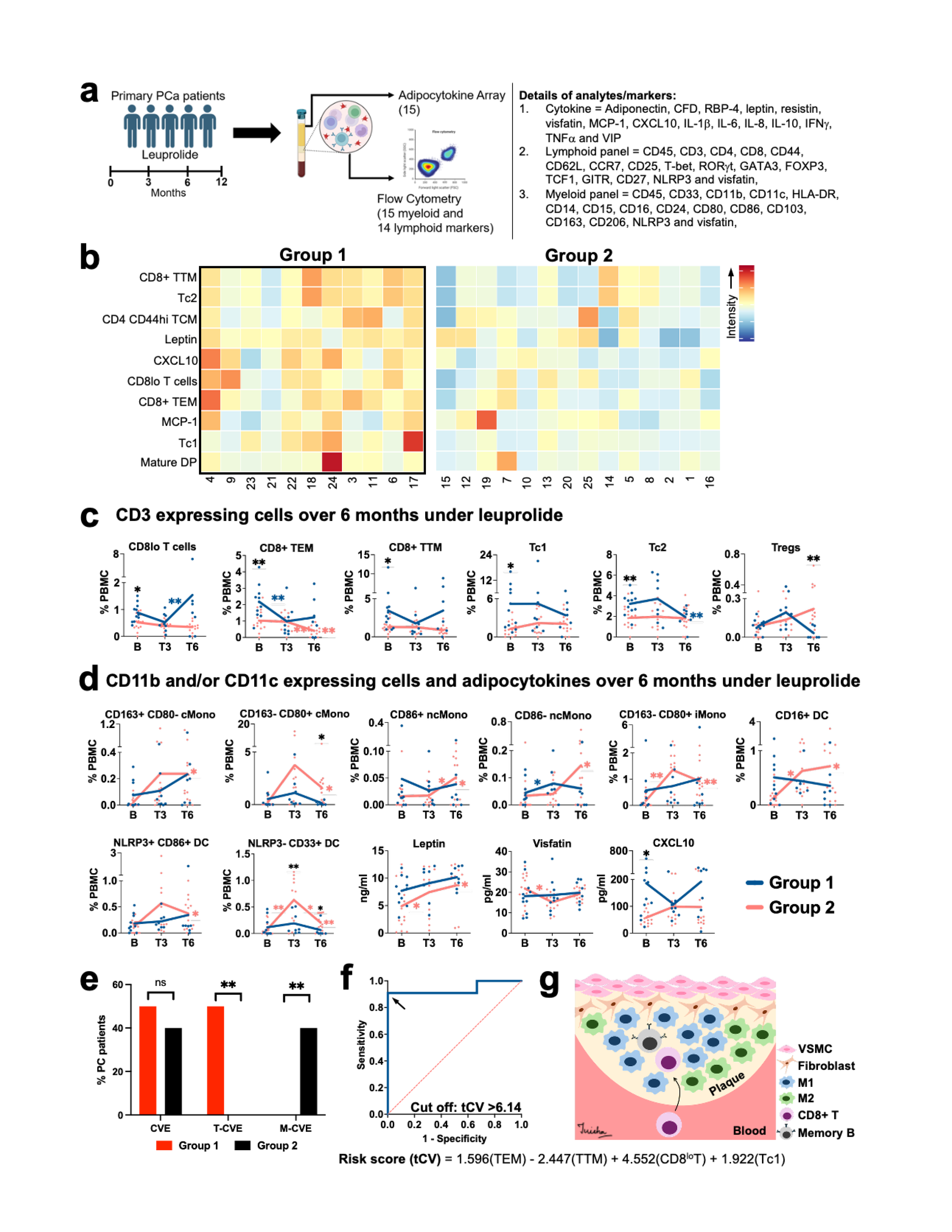
Circulating CD8^+^T cell states identify a phenotype associated with leuprolide-related cardiovascular risk. **a,** Schematic of the CoPro study and longitudinal blood collection used to investigate circulating immune biomarkers associated with cardiovascular events (CVE) during leuprolide treatment in patients with primary prostate cancer (PC). Peripheral blood samples collected at baseline and during treatment were analyzed by multiparametric flow cytometry and adipocytokine profiling. **b,** Unsupervised clustering of baseline circulating immune and soluble parameters separates patients into two major groups, including a T cell-enriched Group 1 and Group 2 lacking this prominent lymphoid phenotype. **c,** Frequencies of indicated circulating T cell populations at baseline (B), and after 3 months (T3) and 6 months (T6) of leuprolide treatment in Groups 1 and 2. **d,** Frequencies of indicated myeloid cell populations and circulating adipocytokine concentrations across baseline, T3 and T6 time points in Groups 1 and 2. Statistical analyses for longitudinal immune and adipocytokine measurements are done using student T-test, presented as *p<0.05 and **p<0.01 (asterisk with black, blue and red colors indicate the comparison of group 1 *vs* group 2, T3 *vs* B or T6 *vs* T3 in group 1 and group 2, respectively) in panel **c** and **d**. **e,** Distribution of retrospectively adjudicated CVE among the immune-defined patient groups. Events were classified as multifactorial cardiovascular events (M-CVE) or treatment-associated cardiovascular events (T-CVE) as detailed in **Supplementary Table 1**. **f,** Receiver operating characteristic (ROC) analysis of the multivariable logistic regression model incorporating Tc1 (cytotoxic type 1 CD8**^+^**T cells), CD8**^+^**effector-memory (TEM), CD8^lo^ and CD8**^+^**transitional-memory (TTM) populations to distinguish the baseline immune-defined patient groups. The corresponding composite tCV score and threshold were derived using the Youden index. **g,** Proposed working model integrating the murine spatial transcriptomic and human circulating immune findings. Leuprolide-associated remodeling of the coronary plaque immune niche involves lymphoid aggregates and inflammatory M1-like macrophages, while a pre-existing circulating CD8**^+^**T cell phenotype identifies patients associated with treatment-related cardiovascular toxicity. Conceptual connections denote relationships requiring direct mechanistic validation. Data were obtained from 25 patients with primary PC enrolled in the CoPro study. The number of evaluable patients at individual longitudinal time points and for cardiovascular outcomes is indicated in the corresponding panels or source data. Quantitative data are presented as mean±s.e.m. Individual patient values are shown where applicable.

Unbiased analysis of the CD3**^+^**lymphoid compartment resolved multiple phenotypically distinct T cell populations (**Supplementary Figure 6a-c**). CD4**^+^**T cells comprised T-bet^hi^Foxp3^lo^GATA3**^-^**GITR**^+^**type 1 helper T cells (Th1), T-bet^lo^Foxp3^hi^GATA3**^+^**GITR**^+^**regulatory T cells (Treg) and T-bet^lo^Foxp3^lo^GATA3**^-^**CCR7**^+^**central memory T cells (TCM), the latter further distinguished by CD44^hi^ and CD44^lo^ phenotypes with concordant CD27 expression patterns. CD8**^+^**T cells resolved into five major states: CCR7**^+^**CD27^lo^GATA3**^-^**CD8^lo^T cells, CCR7**^-^**CD27^lo^GATA3**^-^**transitional memory T cells (TTM), CCR7**^-^**CD27^hi^GATA3**^-^**GITR**^+^**effector memory T cells (TEM), and CCR7**^-^**CD27**^-^**GATA3**^+^**cytotoxic T cells separated into T-bet^hi^type 1 (Tc1) and T-bet^lo^type 2 (Tc2) populations^37,38^. Two additional CD3**^+^**populations were identified: mature CD4**^+^**CD8**^+^**double-positive T cells enriched for GITR and T-bet, and CD4**^-^**CD8**^-^**double-negative T cells. Across the cohort, the overall representation of these lymphoid states remained broadly preserved across baseline and 3-, 6- and 12-month treatment time points (**Supplementary Figure 6d**).

Parallel profiling of the myeloid compartment resolved monocytes, dendritic cells (DC), B cells and natural killer (NK) cells according to CD11b, CD11c, CD33, MHC-II and lineage-associated markers (**Supplementary Figure 7a-c**)^39,40^. Monocytes comprised classical (CD14**^+^**CD16^lo^), intermediate (CD14**^+^**CD16^hi^) and non-classical (CD14**^-^**CD16^hi^) populations, with additional states defined by CD80, CD163 and CD86 expression. DC were further resolved into five phenotypic populations according to CD16, CD103, NLRP3, CD86 and CD33, whereas circulating B cells comprised CD16**^+^** and CD16**^-^**populations. Similar to the lymphoid compartment, the overall distribution of major myeloid populations remained broadly stable across treatment time points (**Supplementary Figure 7d**).

Because non-calcified plaque volume increases only in a subset of leuprolide-treated patients (**Supplementary Figure 1a**), we next examined whether baseline circulating immune heterogeneity could resolve distinct patient populations. Unsupervised clustering integrating the frequencies of 26 immune populations and 15 adipocytokines separated patients into two major groups distinguished predominantly by T cell composition (**Figure 4b**). Approximately 44% of patients comprised a T cell-enriched group (Group 1), characterized particularly by higher proportions of CD8**^+^**T cell subsets, whereas Group 2 lacked this prominent lymphoid phenotype. Within Group 1, CD8^lo^, CD8**^+^**TTM and CD8**^+^**TEM populations transiently decreased after three months of leuprolide treatment but recovered by six months to levels not significantly different from baseline (**Figure 4c**). Tc1, Tc2 and Treg populations showed later trends toward reduction. In contrast, Group 2 exhibited treatment-associated increases in multiple classical, intermediate and non-classical monocyte states, together with CD16**^+^**, NLRP3**^+^**CD86**^+^**and NLRP3**^-^**CD33**^+^**DC populations (**Figure 4d**). Leptin also increased following treatment, reaching significance in Group 2, whereas other circulating immune populations and adipocytokines, including TNFα, IFNγ, IL-1β, IL-6, IL-8, IL-10, adiponectin, adipsin, resistin and visfatin, showed limited changes (**Supplementary Figure 8a, b**). Collectively, these analyses reveal substantial baseline immune heterogeneity among patients with primary PC and identify a stable CD8**^+^**T cell-enriched circulating phenotype in approximately 44% of patients receiving leuprolide, providing a framework to determine whether pre-existing immune states distinguish susceptibility to treatment-associated cardiovascular toxicity.

### Circulating CD8+T cell subsets identify patients with leuprolide-associated cardiovascular risk

We next investigated whether the CD8**^+^**T cell-enriched phenotype was associated with CVE during leuprolide treatment. Events were classified as multifactorial CVE (M-CVE) when attributable to pre-existing cardiovascular disease or established risk factors, or as treatment-associated CVE (T-CVE) when occurring without an evident baseline cardiovascular predisposition (**Supplementary Table 1**). Notably, left ventricular dysfunction classified as T-CVE occurred exclusively among patients in the T cell-enriched Group 1, consistent with the ventricular remodeling observed following leuprolide exposure in mice (**Figure 3a, e**). Although the overall incidence of retrospectively available CVE was comparable between Group 1 and Group 2 (50% *vs* 40%, respectively), T-CVE occurred only in Group 1, whereas events in Group 2 were predominantly classified as M-CVE (**Figure 4e**). We therefore tested whether circulating T cell subsets could collectively distinguish the Group 1 phenotype and potentially leuprolide-associated CVE risk. Multiparametric regression identified Tc1, CD8**^+^**TEM, CD8^lo^T cells and CD8**^+^**TTM as the strongest contributors distinguishing Group 1, which were integrated into a composite leuprolide therapy-associated cardiovascular risk (tCV) score (**Figure 4f**). A tCV threshold >6.14 classified the CVE high-risk Group 1 phenotype with 90% sensitivity and 100% specificity in this discovery cohort. Incorporating additional immune or soluble parameters did not improve model performance, and individual T cell populations were less discriminatory than the composite signature (**Supplementary Figure 9a**). The tCV classification remained informative across longitudinal treatment time points (**Supplementary Figure 9b**), with high-scoring patients displaying visibly greater representation of the contributing CD8**^+^**T cell populations (**Supplementary Figure 9c**). Together, these findings identify a circulating CD8**^+^**T cell signature associated with leuprolide-related cardiovascular toxicity (**Figure 4g**) and nominate the tCV score as a candidate biomarker for prospective validation and treatment-risk stratification.

## Discussion

Cardiovascular toxicity is an increasingly important determinant of long-term adverse outcomes in patients with cancer, particularly in malignancies such as PC, for which effective therapies have substantially extended survival^2,3^. Although cardiovascular complications accompany diverse cancer treatments, their underlying mechanisms vary considerably. Anthracyclines can directly injure cardiomyocytes through oxidative stress and mitochondrial dysfunction^41^, whereas immune-based therapies, including immune checkpoint inhibitors and chimeric antigen receptor T cell therapies, can trigger inflammatory cardiovascular toxicities through dysregulated immune activation^42,43^. ADT has been a cornerstone of PC treatment for decades, yet the biological mechanisms underlying its cardiovascular toxicity remain incompletely understood. Here, by integrating spatially resolved cardiac transcriptomics with longitudinal peripheral immune profiling in patients, we identify vascular immune remodeling as a potential mechanism underlying leuprolide-associated cardiovascular toxicity. Our findings further reveal a circulating CD8**^+^**T cell phenotype associated with treatment-related CVE, providing a potential framework for identifying patients particularly susceptible to GnRH agonist-associated cardiovascular complications.

Coronary atherosclerosis represents a major substrate for CVE in patients receiving ADT^5,6^. Coronary computed tomography angiography in the Revolution-1 study demonstrated increased non-calcified plaque volume following ≥ 6 months of leuprolide treatment, whereas relugolix did not produce a comparable increase^5^. This distinction is particularly relevant because both therapies effectively suppress circulating testosterone, suggesting that differential cardiovascular effects cannot be attributed solely to systemic androgen deprivation. Clinical studies similarly suggest a higher incidence of major CVE with GnRH agonism than antagonism^4^. Experimental studies have implicated GnRH agonism in macrophage recruitment and expansion of necrotic plaque regions, while treatment-associated follicle-stimulating hormone surges may further enhance vascular inflammation through VCAM1-dependent recruitment of inflammatory cells^44–46^. Our findings extend these observations by resolving the cellular architecture of the leuprolide-exposed coronary plaque and identifying plaque fibroblasts, lymphoid aggregates and inflammatory macrophages as components of a coordinated vascular immune response.

A particularly notable finding was the compartment-specific response of fibroblasts to leuprolide. Inferred GnRHR-signaling activity increased in plaque-resident fibroblasts while decreasing in fibroblasts within histologically normal vasculature, suggesting that transcriptional responses to GnRH agonism are strongly influenced by the local tissue state. Increased AR-signaling activity was also observed specifically in plaque fibroblasts despite systemic androgen deprivation. These alterations coincided with enrichment of T+B cell lymphoid aggregates and M1-like macrophages and with extensive remodeling of predicted intercellular communication. Leuprolide attenuated fibroblast-derived *Fn1* and *Bgn* interactions associated with extracellular matrix organization while increasing inflammatory *C3*- and *Cd74*-associated programs. T+B lymphoid aggregates exhibited IRF1- and CIITA-associated regulatory activity, whereas plaque M1-like macrophages acquired IFNα- and IFNγ-response programs and persistent mesenchymal-remodeling signatures. Together, these observations support a model in which GnRH agonism reorganizes an existing inflammatory plaque niche rather than uniformly activating the cardiovascular system. Importantly, normal vasculature exhibited a distinct M2-like macrophage- and complement-associated response, emphasizing that plaque and normal vasculature respond differently to the same systemic therapy.

The clinical immune profiling provides an important translational counterpart to this spatially resolved biology. Approximately 44% of patients displayed a pre-existing CD8**^+^**T cell-enriched circulating phenotype characterized predominantly by increased CD8**^+^**T cell subsets. Although overall CVE frequencies were similar between immune-defined patient groups, events without an evident baseline cardiovascular predisposition occurred selectively within the T cell-enriched group. A composite signature incorporating Tc1, CD8**^+^**effector memory, CD8**^+^**transitional memory and CD8^lo^T cell populations distinguished this phenotype more effectively than individual immune populations. These observations suggest that the host immune state before treatment may influence susceptibility to cardiovascular toxicity following GnRH agonism. The resulting tCV score therefore represents a candidate biomarker for prospective evaluation rather than an established clinical predictor. If independently validated, such an approach could complement conventional cardiovascular risk assessment when selecting between GnRH agonists and antagonists.

Our spatial analyses also identified cardiac alterations beyond the plaque. Leuprolide was associated with an approximately 4-fold expansion of hypertrophic ventricular cardiomyocytes accompanied by oxidative phosphorylation and myogenesis programs. Notably, these hypertrophic cells lacked detectable AR or GnRHR activation and showed limited predicted communication with other cardiomyocyte populations. These findings argue against a simple cell-autonomous effect of leuprolide on hypertrophic cardiomyocytes and raise the possibility that ventricular remodeling may develop secondary to vascular or tissue-level perturbations^35,36^. Consistent with this observation, ventricular dysfunction occurred among patients exhibiting the CD8**^+^**T cell-enriched phenotype. Establishing a causal relationship between plaque inflammation and ventricular remodeling will, however, require longitudinal functional cardiac studies.

A second tissue program emerged within cardiac adipose and stromal compartments. Leuprolide reduced AR and GnRHR transcriptional activities in adipocytes and was associated with increased adipocyte-memory B cell communication, including *C3*- and *Lpl-Lrp1*-associated interactions. Adipocytes and memory B cells shifted from metabolic programs toward mesenchymal-remodeling states, while myofibroblast accumulation and altered stromal-immune communication were evident in adjacent cardiac compartments. Circulating leptin increased during treatment, particularly among patients without the baseline T cell-enriched phenotype. These observations suggest that adipose-stromal remodeling may represent a mechanistically distinct component of ADT-associated cardiovascular toxicity, complementing the inflammatory remodeling observed within vulnerable calcified plaque of patients^6^.

Several limitations warrant consideration. Spatial transcriptomic analyses were performed in a small number of biological replicates and should therefore be interpreted as discovery level evidence requiring orthogonal validation. The high-fat diet/resiquimod in murine PC model establishes pre-existing vascular inflammation but may not fully reproduce human atherosclerosis. Predicted ligand-receptor interactions and transcriptional pathway activities do not establish causal signaling, and the mechanisms producing opposing GnRHR responses in plaque fibroblasts and adipocytes remain unresolved. Most importantly, the clinical cohort was small, and cardiovascular outcome information was available for only a subset of participants. The tCV signature therefore requires validation in larger, independent and prospectively phenotype cohorts before clinical implementation.

Collectively, our findings support a model in which leuprolide orchestrates distinct cardiovascular immune niches, coupling inflammatory lymphoid-macrophage organization within vulnerable plaque with adipose-stromal and cardiac remodeling. The identification of a circulating CD8**^+^**T cell signature associated with treatment-related CVE further connects localized vascular immune biology with systemic host immunity. These findings provide a framework for understanding differential cardiovascular toxicity among ADT modalities and suggest that immune-informed risk stratification could ultimately contribute to individualized selection of hormonal therapy in patients with PC.

## Methods

### Coronary plaque analysis

Coronary plaque progression in PC patients was assessed based on data available from the REVELUTION-1 (NCT05320406), an open-label study of men with nonmetastatic PC conducted across four Emory University-affiliated centers in Atlanta, Georgia. Trial was approved by the Emory University Institutional Review Board, conducted in accordance with the Declaration of Helsinki, and all participants provided written informed consent. The complete trial design, eligibility criteria, treatment procedures, coronary computed tomography angiography (CCTA) acquisition and plaque quantification methodology have been reported previously^5^. For the analysis presented in Supplementary Figure 1a, data were extracted from 57 evaluable patients receiving radiation therapy alone (U means no drug-treatment) or radiation therapy plus leuprolide (L) with available baseline and 12-month CCTA measurements. Coronary plaque burden was quantified as previously described and included total plaque volume (TPV), non-calcified plaque volume (NCPV), calcified plaque volume (CPV) and low-attenuation plaque volume (LAPV). For each patient, the absolute change in each plaque measure was calculated from baseline to the 12-month assessment and compared between treatment groups. These data were used to define the leuprolide-associated coronary plaque phenotype.

### Murine prostate cancer model and leuprolide treatment

To investigate the mechanisms underlying leuprolide-associated cardiovascular complications, we established a murine model combining PC and atherogenic inflammation. FVB/NJ male mice (Jackson laboratory, 001800, 8 weeks old) were maintained under pathogen-free conditions with *ad libitum* access to food and water in accordance with protocols approved by the Institutional Animal Care and Use Committee (IACUC) at Emory University. High-fat diet (Envigo, TD.88317 adjusted calorie diet) was given for 8 weeks before tumor implantation and continuing throughout the experiment. Toll-like receptor 7/8 agonist, resiquimod (3.4 mg/kg in 1:3 of ethanol:acetone mixture, topical application on ear, 3 times a week; total three weeks; Selleckchem, S8133) administration was initiated with high-fat diet. This combined approach was selected to promote baseline vascular inflammation and facilitate assessment of treatment-associated alterations in coronary plaque immune composition.

Next, animals were randomly assigned to untreated (subcutaneously received 40 μL phosphate buffer saline, PBS) or leuprolide (0.47 mg/kg, subcutaneously; Sigma-Aldrich, L0399) treatment groups and inoculated with Myc-CAP cell line (1x10^6^ cells, subcutaneously; ATCC, CRL-3255). Tumor kinetics monitored weekly using Vernier calipers, and treatment started once tumor reached approximately 200 mm^3^ size within 3 weeks period. At peri-castration stage, while tumors were responding to leuprolide treatment (approximately 10 days from initiation), mice were euthanized, whole heart harvested and formalin-fixed paraffin embedded blocks (FFPE) were prepared.

### Spatial compartment delineation

Serial sectioning (5μm) and hematoxylin and eosin (H&E) staining were done on cardiac blocks until coronary plaque was identified by an experienced cardiovascular pathologist who was blinded to treatment allocation throughout the study. These sections from each heart were allocated for spatial transcriptomic profiling. Histologically, coronary plaque was identified based on intimal thickening, inflammatory-cell accumulation, disruption of normal vascular architecture, necrotic and lipid-rich vascular lesions, whereas vessels lacking plaque formation were annotated as normal vasculature. In addition, cardiac adipose tissue, atrial myocardium and ventricular myocardium were independently identified based on anatomical and histological features. These annotations generated five predefined anatomical compartments for downstream analyses: coronary plaque (P), normal vasculature (Va), adipose tissue (Ad), atria (A) and ventricle (V).

### Visium HD spatial transcriptomic library preparation and sequencing

Emory Integrated Genomics Core performed spatial transcriptomic profiling of cardiac sections prepared from untreated and leuprolide-treated mice using the Visium HD Spatial Gene Expression platform (10x Genomics). RNA was assessed using an Agilent Bioanalyzer to confirm suitability of selected cardiac section for probe-based spatial gene-expression analysis. Spatial gene expression libraries were generated using the Mouse Transcriptome, 6.5-mm kit (10x Genomics, PN 1000676; 4 reactions) together with the Visium CytAssist workflow. Briefly, mouse whole-transcriptome probe pairs were hybridized to tissue RNA and cognate probe pairs were ligated *in situ*. Tissue sections were positioned for CytAssist processing such that the region of interest was contained within the 6.5-mm capture area. Ligated gene-expression probes were released from the tissue and transferred onto the Visium HD slide, where they were captured by spatially barcoded oligonucleotides with continuous array of 2x2 µm squares within the 6.5 mm capture area, enabling spatially resolved expression profiling without tissue dissociation^1^. Following this, Visium HD slides were removed from the CytAssist instrument and processed for gene expression library construction. Spatially barcoded libraries were amplified and indexed using the Dual Index Kit TS Set A (10x Genomics, PN 1000251). Library concentration and quality were assessed before sequencing. Indexed libraries were pooled and sequenced using an Illumina NextSeq 2000 platform with the recommended reagents (Illumina, PN 20100990). The resulting sequencing data retained spatial barcode and transcript information required to reconstruct spatial transcriptomic maps for each cardiac section.

### Sequencing data preprocessing, cluster annotation and regional analyses

Raw sequencing data were processed using Space Ranger v3.1.1 (10x Genomics) with mouse transcriptome probe set v2.0 and mm10-2020-A reference for sequencing-read processing, spatial barcode assignment, probe filtering, image alignment and generation of spatial gene-expression matrices. Downstream analyses were performed using the 8 µm spatial bin output as it increased counts per analytical unit while preserving the tissue context required for compartment-resolved cardiac analyses. Spatial gene expression matrices and corresponding spatial coordinates were imported into Scanpy v1.11.4^2^ for quality control on each cardiac section. Quality-control thresholds were calculated separately for each specimen from the 5th and 95th percentiles of the observed distributions. Bins were retained when detected genes were between max(10, sample-specific 5th percentile) and the sample-specific 95th percentile, total counts were between max (20, sample-specific 5th percentile) and the sample-specific 95th percentile, mitochondrial-transcript percentage was no greater than min (20%, sample-specific 95th percentile), and log10(detected genes)/log10(total counts) was at least max (0.8, sample-specific 5th percentile). No additional gene level minimum-bin filter was applied after concatenation. Ultimately, 700,410 of 1,363,154 spatial bins were retained across the four cardiac specimens. Gene expression counts were normalized to 10,000 counts per spatial bin and log-transformed using log1p. The 3,000 most highly variable genes were identified using Scanpy Seurat dispersion method, with specimen identity used as the batch key.

Spatially informed dimensionality reduction and clustering were performed using BANKSY v1.3.4^3^, incorporating both transcriptional expression and local spatial neighborhood information. BANKSY features were computed using 15 spatial neighbours, scaled-Gaussian distance weighting, a neighbourhood expression weight of λ = 0.2 and maximum azimuthal harmonic order m = 1. Principal component analysis (PCA) was performed using 20 components, and sample-associated variation was corrected using Harmony^4^, while retaining specimen identity for subsequent analyses. A nearest-neighbor graph was constructed from the Harmony-adjusted BANKSY representation using 20 neighbors, and Uniform Manifold Approximation and Projection (UMAP) was used to visualize transcriptional relationships among spatial profiles. Unsupervised clustering was performed using the Leiden algorithm at a resolution of 0.6, resulting in 50 transcriptionally distinct clusters across the four cardiac specimens. Cellular identities were assigned to transcriptional clusters based on differentially expressed marker genes, established lineage-associated markers and their spatial localization within corresponding histological structures. Marker genes for each cluster were identified using the two-sided Wilcoxon rank-sum test. Cluster identities were subsequently overlaid onto the spatial transcriptomic maps to evaluate their anatomical distribution across cardiac tissues. Further, polygon coordinates exported from the pathologist-defined regions (P, Va, Ad, A and V) were mapped to spatial-transcriptomic coordinates using the sample-specific high-resolution image scale factor, and a point-in-polygon operation assigned each 8 µm bin to pathologist-defined regions. Compartment-focused functional analyses were restricted to biologically relevant BANKSY clusters selected according to marker expression and spatial localization. The cluster sets used for each region and analysis are specified in the corresponding figure. All 50 clusters were retained in the complete dataset and global and region-specific visualizations. Further, each annotated population was quantified as the number of bins assigned to that cluster and expressed as a proportion of the total annotated bins within the corresponding compartment or cardiac section. Because the observations were 8 µm bins rather than segmented cells, cluster labels were interpreted as putative cell populations or states represented by spatial profiles.

### AR and GnRHR signaling activity analysis

AR- and GnRHR-associated transcriptional activities were inferred from log-normalized expression using the univariate linear model (ULM) in decoupler v2.1.4^5^. Inferred AR-associated activity was assessed using the NELSON_RESPONSE_TO_ANDROGEN_UP gene signature, whereas the KEGG_GNRH_SIGNALING_PATHWAY gene signature was used to assess inferred GnRHR-signaling activity. Higher scores may indirectly reflect downstream inferred AR- or GnRHR-signaling but do not constitute direct measurements of receptor activation. Resulting activity scores were integrated with the previously defined cluster identities and compared between untreated and leuprolide-treated groups across pathologist-annotated anatomical compartments. For the figure panels, profiles were restricted to ULM scores greater than zero before plotting and testing. Within each region, groups were compared within clusters using two-sided Mann-Whitney U tests, followed by Benjamini-Hochberg correction across clusters.

### Pathway analysis

Biological programs were characterized for predefined cellular populations present in the anatomical compartments of both groups using Hallmark gene set over-representation analysis (ORA). Gene expression was first compared between leuprolide-treated and untreated profiles using the Wilcoxon rank-sum test implemented in Scanpy, with up to 500 genes returned in each direction. Genes were ranked using a signed composite score = sign (log2 fold change) x -log10 (Benjamini-Hochberg-adjusted P value), and the 200 genes with the largest absolute scores across both directions were separated into leuprolide-high and untreated-high query sets. Fifty mouse Hallmark gene sets obtained from the Molecular Signatures Database (MSigDB)^6^ were tested using one-sided Fisher exact tests. The background comprised genes present in the corresponding bidirectional differential expression export. Statistical significances were adjusted across Hallmark sets within each region-cluster-direction analysis using the Benjamini-Hochberg method.

### Ligand-receptor interaction and cellular communication analysis

Cellular communications were inferred for pathologist-defined anatomical compartments using ChatSpatial v1.2.1^7^ in combination with LIANA v1.7.1^8^ and the ‘mouseconsensus’ ligand-receptor resource. Cellular populations represented by fewer than 10 spatial profiles were excluded from communication analysis. Ligand-receptor inference required a minimum non-zero expression proportion of 0.01 and used 1,000 permutations. Communication networks were inferred separately after pooling profiles from untreated or leuprolide-treated specimens. For differential visualization, untreated and treated results were outer-joined by sender, receiver, ligand and receptor; combinations absent from one condition were assigned an ‘lr_means’ value of zero, and the untreated value was subtracted from the treated value. Heatmaps show the sum of these differences for each sender-receiver pair, whereas circos plots show the 25 pairs with the largest absolute summed differences. Directionality was assigned according to ligand expression in the predicted sender population and cognate receptor expression in the predicted receiver population. Differential communication patterns were visualized using heatmaps and circos plots, with edges representing the direction and relative change in predicted ligand-receptor communication between leuprolide-treated and untreated conditions. Although anatomical regions were defined from spatially resolved transcriptomic data, physical distance or spatial adjacency between sender and receiver profiles was not incorporated into ligand-receptor inference. Thus, inferred networks represent descriptive, transcriptionally supported communication among cellular populations residing within the same pathologist-defined anatomical compartment rather than evidence of direct physical cell-cell contact.

### Transcription factor activity

Transcription factor (TF) activities were inferred from transcriptomic profiles using the ULM in decoupler v2.1.6 with the mouse CollecTRI regulon^9^, comprising 43,226 signed TF-target gene interactions. TF represented by at least five target genes were retained. The directionality of individual regulatory interactions was retained, enabling coordinated expression of positively and negatively regulated target genes to contribute to inferred TF activity. Log-normalized gene expression matrices from the 8 µm spatial bin were used as input. For each transcription factor, ULM modeled the expression of its target genes according to the signed regulatory relationships contained within the CollecTRI network and generated an activity score representing the relative transcriptional influence of that regulator within each bin. Exploratory treated *vs* untreated comparisons used two-sided Mann-Whitney U tests across spatial profiles, with Benjamini-Hochberg correction across TF within each region-cluster comparison. Because spatial profiles were nested within mice, these tests were not interpreted as replicate-level treatment inference. For each region-cluster comparison, the displayed bars show the difference in mean profile-level activity between leuprolide-treated and untreated groups for the 20 TF with the largest absolute differences, bar selection was based on effect magnitude rather than statistical significance. These scores estimate relative regulon-consistent activity.

### Cardio-Oncology Prostate cancer (CoPro) biobank

Serial blood specimens were obtained from 25 primary PC patients enrolled in the prospective CoPro study during 3/2021-4/2023 at Emory University-affiliated Hospitals, as per approved Institutional Review Board (00002057) and the Declaration of Helsinki. All participants provided written informed consent. Blood collected at baseline (B) as well as following 3, 6 and 12 months of leuprolide treatment (T3, T6 and T12, respectively). Specimens were centrifuged 500g for 10 minutes and plasma were collected. Peripheral blood mononuclear cells (PBMC) were further isolated from remaining pellet using Ficoll-based centrifugation method^47^. These samples were stored in liquid nitrogen until immune cell-profiling and adipocytokine assay were conducted.

Clinical CVE information was retrospectively available for 14 of the 25 patients through the review of clinical records and available in Supplementary Table 1. CVE were further categorized according to whether the event occurred in the context of pre-existing cardiovascular disease or established cardiovascular risk factors, considered multifactorial cardiovascular events (M-CVE). Events occurring during leuprolide treatment that were considered unexpected based on documented cardiovascular history and baseline risk factors, were designated treatment-associated cardiovascular events (T-CVE). Two independent cardiologists had utilized this stratification to define T-CVE *vs* M-CVE in each patient, and their collective outcome was considered in Supplementary Table 1.

### High-dimensional flow cytometry-based immunophenotyping

Cryopreserved PBMC were thawed, washed and resuspended in PBS for staining. The lymphoid panel included antibodies against CD45 (Biolegend, 304026), CD3 (Biolegend, 300448), CD4 (ThermoFisher scientific, 15-0049-42), CD8 (Biolegend, 344712), CD44 (Biolegend, 103049), CD62L (ThermoFisher scientific, 368-0629-42), CCR7 (Biolegend, 353264), CD25 (Biolegend, 302614), T-bet (Fisher scientific, BDB568167), RORγt (ThermoFisher scientific, 46-6988-82), GATA3 (Biolegend, 653812), FOXP3 (Biolegend, 320126), TCF1 (Biolegend, 655208), GITR (Biolegend, 371212), CD27 (ThermoFisher scientific, 366-0279-42), NLRP3 (Bio-techne, IC7578A) and visfatin (R&D systems, IC4044V), together with a fixable viability dye. The myeloid panel included antibodies against CD45, CD33 (BD Biosciences, 741612), CD11b (Biolegend, 301408), CD11c (Biolegend, 741827), HLA-DR (BD Biosciences, 561224), CD14 (ThermoFisher scientific, MHCD1418), CD15 (Biolegend, 301917), CD16 (ThermoFisher scientific, H006T03B06), CD24 (Biolegend, 311120), CD80 (Biolegend, 305225), CD86 (Biolegend, 305440), CD103 (BD Biosciences, 748503), CD163 (Biolegend, 333646), CD206 (Biolegend, 321142), NLRP3 and visfatin, again together with a fixable viability dye. For surface immunophenotyping, PBMC were incubated with the corresponding surface-antibody cocktail and fixable viability dye for 30 min at room temperature. Cells were subsequently washed with PBS, subjected to fixation and permeabilization with recommended reagents as per manufacturer protocol (Biolegend, 421403). Intracellular and intranuclear markers, including T-bet, RORγt, GATA3, FOXP3, TCF1 NLRP3 and visfatin were stained in permeabilization buffer according to the optimized staining protocol of manufacturer (Biolegend, 421403). Following completion of staining, cells were washed, resuspended in PBS and data acquired using Cytek Aurora 5L.

Flow cytometry data were analyzed using FlowJo v10.7, with help of recommended tutorial. Initial quality control was performed using the FlowAI together with manual inspection of individual files and corresponding fluorescence-minus-one (FMO)/isotype controls to identify and exclude poor-quality samples or events. Following this, debris, doublets and non-viable events were excluded. For lymphoid analysis, viable CD3**^+^**T cells were selected, whereas myeloid analysis included viable cells expressing CD11b, CD11c or both markers. These cellular populations were concatenated separately for the lymphoid and myeloid datasets while retaining patient identity and collection time point for subsequent sample-level quantification. Unsupervised clustering was performed on concatenated dataset using the FlowSOM plugin. Relationships among phenotypically related populations were assessed using Euclidean-distance-based analysis, and UMAP was used for dimensionality reduction and visualization of the resulting high-dimensional immune-cell populations. ClusterExplorer and HyperFinder were subsequently used together with individual marker expression patterns to characterize and assign biological identities to the FlowSOM-defined clusters. Following cluster annotation, the frequency of each immune cell was determined with respect to each individual total live PBMC.

### Adipocytokine assay

Plasma were thawed and analyzed for adipocytokines (Adiponectin, CFD, RBP-4, leptin, resistin, visfatin, MCP-1, CXCL10, IL-1β, IL-6, IL-8, IL-10, IFNγ, TNFα and VIP) as per manufacturer protocol (Biolegend, 741062; ThermoFisher, EH482RB; and R&D systems, NBP2-82390).

### Patient stratification and tCV score

Baseline patient clustering used 25 participants and 41 variables, comprising 26 flow cytometry population frequencies and 15 soluble mediator concentrations. Soluble-mediator values were transformed using log1p, whereas flow cytometry frequencies were not transformed. Variables were standardized to zero mean and unit variance, after which missing standardized values were replaced by the corresponding variable median. K-means clustering was evaluated for k = 2-6 using Euclidean distance and 10 initializations. For heatmap visualization, variables and patients were ordered independently using Ward hierarchical clustering of Euclidean distances. A contiguous block of 11 patients in the ordered heatmap was designated post hoc as one group, with the remaining 14 patients assigned to the second group. To determine whether the immune features distinguishing these groups were associated with leuprolide-associated cardiovascular outcomes, candidate immune populations were evaluated individually and subsequently incorporated into a multivariable logistic regression model to distinguish the two patient groups. The resulting composite signature was designated the tCV score.

### Statistical analysis

Python v3.10.16 was used for Visium HD preprocessing and quality control, BANKSY clustering, regional annotation, signature scoring and baseline clinical clustering and silhouette analysis. Python v3.12.0 was used for ChatSpatial/LIANA ligand-receptor analysis, differential-expression and Hallmark ORA workflows, and CollecTRI/ULM transcription-factor activity analysis. Additional statistical analyses were performed using GraphPad Prism v11.0.2 in accordance to the above described analysis-specific procedures. No statistical method was used to predetermine sample size. For murine experiments, 4 mice were randomly assigned to treatment groups, and tumor-ulcerated 2 mice were excluded. Summary statistics and distributional displays are defined in the corresponding figure legends, and categorical variables are presented as counts or percentages.

For murine spatial transcriptomic experiments, individual mice were considered the biological replicates, whereas 8 µm bins represented spatially resolved observations nested within individual specimens. A total of four hearts (two untreated and two leuprolide-treated mice) were analyzed. Accordingly, bin-level analyses were used to characterize spatially resolved transcriptional states and generate mechanistic hypotheses and were not considered independent biological replication. Statistical tests used for individual comparisons are specified above as well as briefly in the corresponding figure legends.

For clinical analyses, the individual patient was considered the biological unit. Longitudinal immune cell frequencies and adipocytokine amounts were analyzed according to treatment time point and immune-defined patient group. The CoPro biomarker cohort comprised 25 patients, with retrospective CVE information available for 14 patients. Individual patient values are shown where appropriate to illustrate interpatient variability. CVE frequencies were compared between patient groups using categorical Chi-square test and also specified in the corresponding figure legends. For development of the tCV score, individual candidate immune cell populations were first evaluated using univariable logistic regression, whereas the composite model was generated using multivariable logistic regression. Model discrimination was evaluated using receiver operating characteristic (ROC) analysis and area under the ROC curve (AUC), and the optimal classification threshold was determined using the Youden index. Sensitivity and specificity were calculated at the selected threshold. For the Revelution-1 analysis, absolute changes in TPV, NCPV, CPV and LAPV from baseline to 12 months were compared between the radiation-alone and radiation+leuprolide groups using a two-sided unpaired Student t-test.

All statistical tests were two-sided, unless otherwise specified. Statistical significance was defined as p<0.05, p<0.01, p<0.001 and p<0.0001 after multiple-testing corrections where applicable. Sample sizes, biological replicates and statistical tests applicable to individual experiments are provided in the corresponding figure legends and source data. Given the exploratory nature and limited size of the CoPro cohort, analyses relating circulating immune phenotypes and the tCV score to cardiovascular outcomes were considered hypothesis-generating and require validation in an independent prospective cohort.

## Data availability

Raw and processed spatial transcriptomic sequencing data will be deposited in the Gene Expression Omnibus (GEO) and publicly available upon publication. Data supporting the findings of this study are available from the corresponding author upon reasonable request.

## Code availability

Spatial profilings were performed using established computational packages, including Scanpy, BANKSY, Harmony, decoupler, ChatSpatial and LIANA, with custom scripts used for preprocessing, regional analyses, pathway and signaling inference, transcription factor activity, cellular communication and visualization. High-dimensional flow cytometry analyses were performed using FlowJo and associated plugins. Custom code required to reproduce the principal analyses and figures will be made publicly available through GitHub upon publication.

---

**Supplementary Figure 1.**
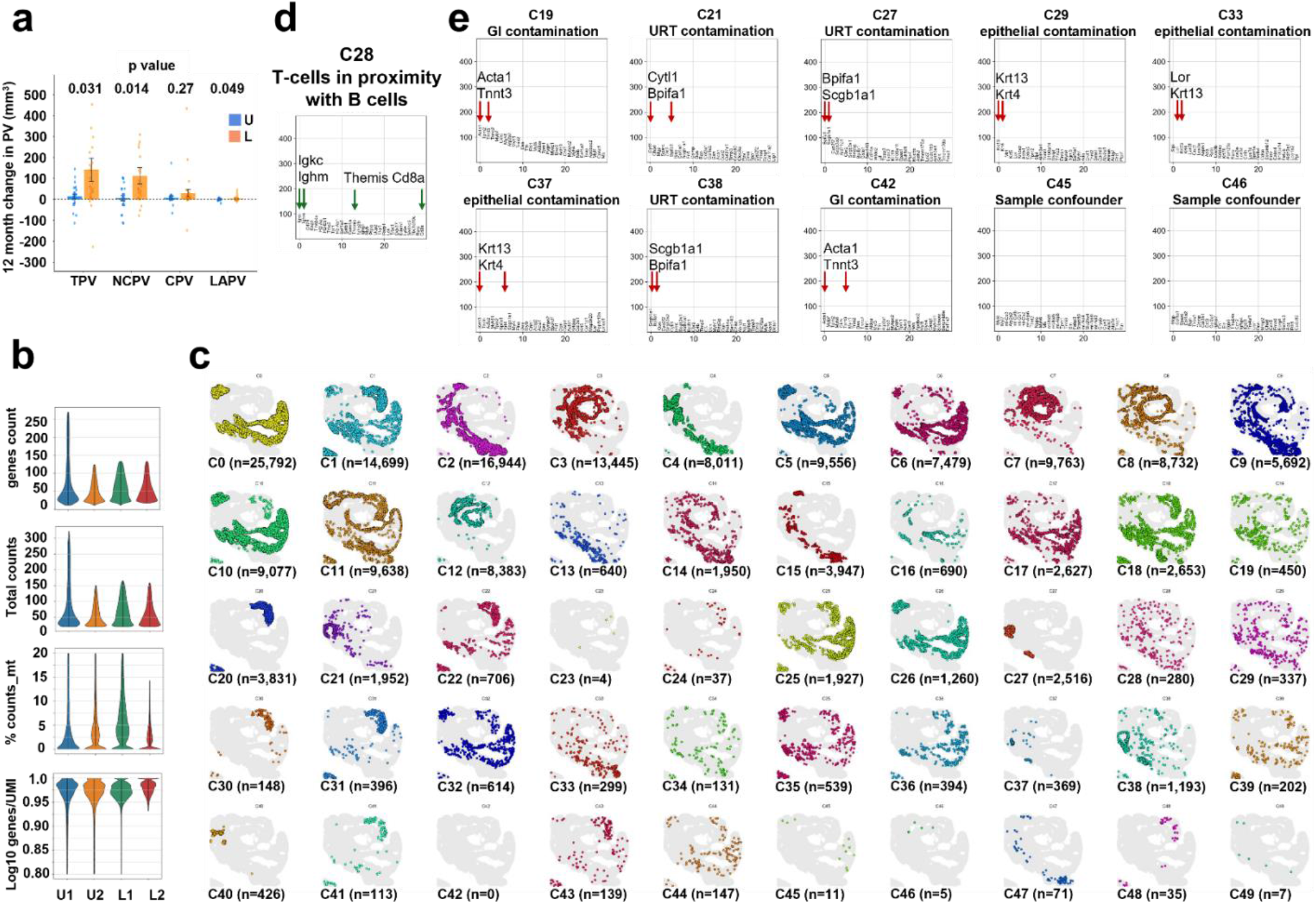
Clinical and spatial transcriptional features of coronary plaque and cardiac tissues. **a,** Absolute changes in total plaque volume (TPV), non-calcified plaque volume (NCPV), calcified plaque volume (CPV) and low-attenuation plaque volume (LAPV) from baseline to 1 year in patients with prostate cancer enrolled during Revolution-1 study for 1) control arm means radiation therapy alone (U) or 2) treatment arm means radiation therapy plus leuprolide (L). **b,** Post-filtering quality-control distributions for spatial transcriptomic datasets generated from untreated (U1 and U2) and leuprolide-treated (L1 and L2) murine hearts, showing detected genes, total counts, mitochondrial-transcript percentage and library-complexity metric by sample. **c,** Spatial maps of the 50 transcriptionally defined cellular populations annotated in **Figure 1c,d**. Map represents sample L2. **d,** Ranked differentially expressed marker genes associated with the T+B cell-enriched lymphoid aggregate population, supporting annotation of spatial regions containing closely associated T cell and B cell transcriptional signatures. **e,** Ranked differentially expressed genes defining sample-restricted populations and tissue contaminants identified during unbiased clustering. These populations, including gastrointestinal- and respiratory-associated transcriptional signatures introduced during tissue collection, were excluded from downstream cardiac compartment analyses. Panel **a** includes 57 patients with primary prostate cancer; data are presented as mean±s.e.m. Panels **b-e** comprise 700,410 post-QC spatial profiles from four murine hearts (two untreated and two leuprolide-treated). U1 and U2 denote individual untreated mice; L1 and L2 denote individual leuprolide-treated mice.

**Supplementary Figure 2.**
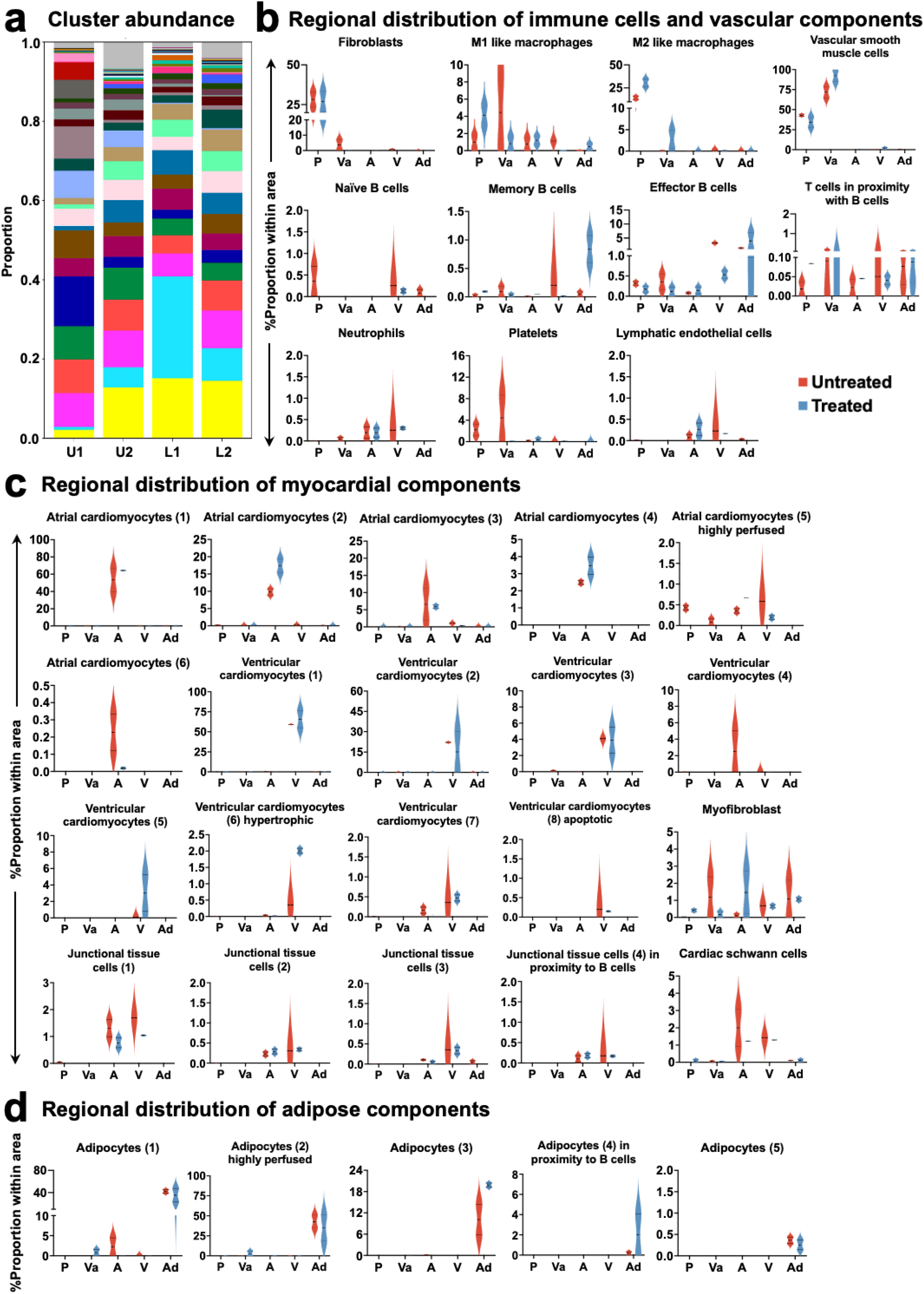
Cellular composition across cardiac tissue compartments following leuprolide exposure. **a,** Relative abundance of the transcriptionally defined populations identified in **Figure 1c,d** across whole-heart sections from individual untreated (U1 and U2) and leuprolide-treated (L1 and L2) mice. **b-d,** Compartment-resolved abundance of cellular populations within pathologist-delineated coronary plaque (P), normal vasculature (Va), cardiac adipose tissue (Ad), atria (A) and ventricle (V), as defined in **Figure 1f**. **b,** Relative abundance of immune and vascular populations, including macrophage, lymphoid, fibroblast and vascular smooth muscle cell subsets, across the indicated anatomical compartments. **c,** Relative abundance of myocardial populations, including atrial and ventricular cardiomyocyte states, junctional tissue and myofibroblast populations. **d,** Relative abundance of adipocyte populations across the indicated anatomical compartments in untreated and leuprolide-treated hearts. Analyses comprised 33,026, 3,830, 12,229, 19,612 and 16,521 spatially resolved cellular profiles within plaque, normal vasculature, adipose tissue, atrial and ventricular regions, respectively, obtained from four murine hearts (two untreated and two leuprolide-treated). Data represents as mean±s.e.m. Statistical analyses were done using student T-test (panel **b, c, d**).

**Supplementary Figure 3.**
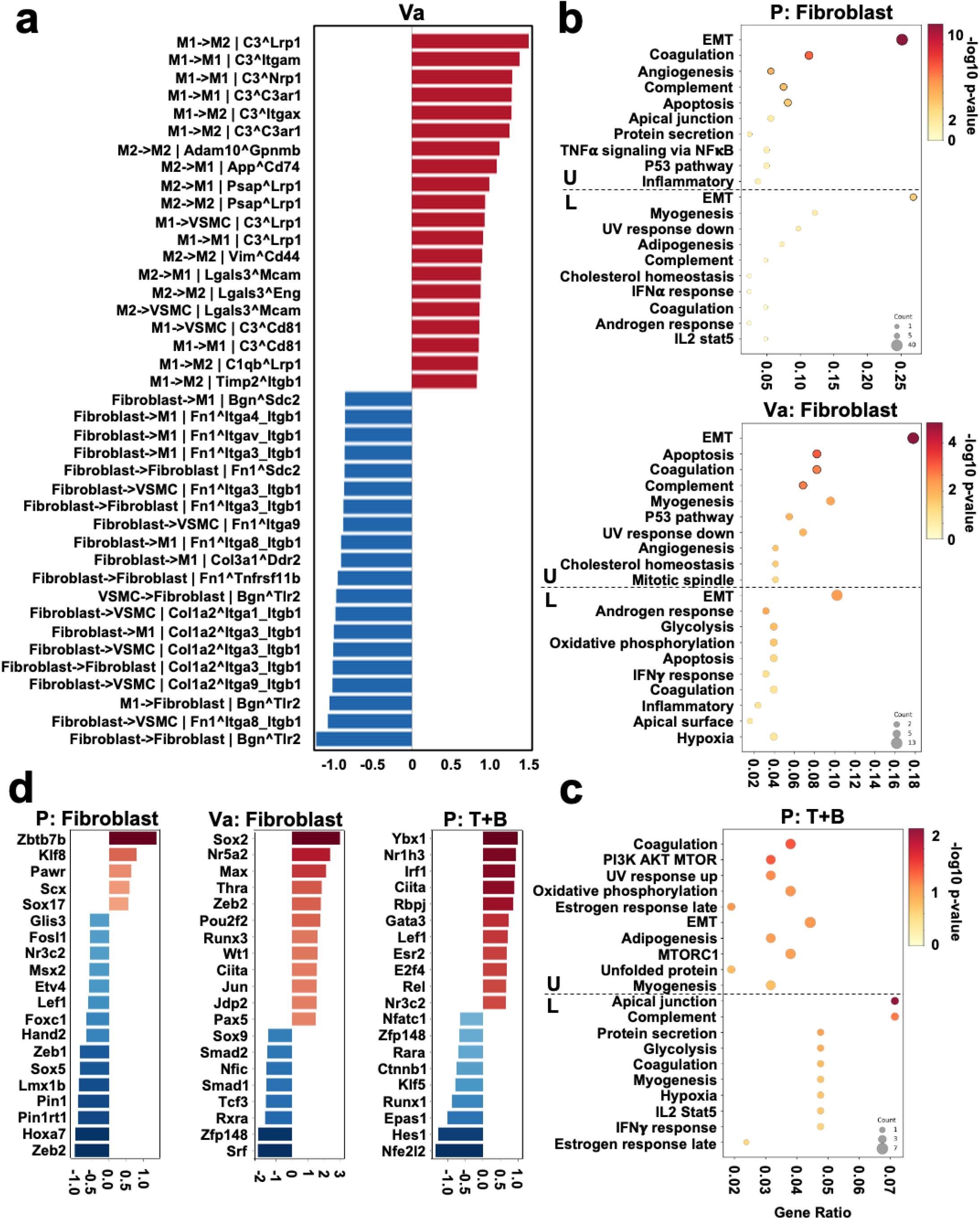
Leuprolide remodels fibroblast and lymphoid transcriptional programs within coronary plaque. **a,** Ligand-receptor interactions with the largest descriptive differences among cellular populations within histologically normal vasculature (Va) following leuprolide treatment relative to untreated controls. Experimental design is shown in **Figure 1a**. Interactions were inferred from spatial transcriptomic expression of cognate ligand-receptor pairs. Arrows indicate signaling direction from ligand-expressing sender to receptor-expressing receiver populations. Bars show the treated-minus-untreated difference in LIANA consensus magnitude for the top 20 leuprolide-high (red color) and top 20 untreated-high (blue color) sender-receiver ligand-receptor combinations. **b,c,** Hallmark pathway enrichment analysis of fibroblasts (**b**) and T+B cell-enriched lymphoid aggregates (**c**) present in coronary plaque (P) and normal vasculature (Va) of untreated (U) and leuprolide-treated (L) hearts. **d,** CollecTRI/ULM transcription factor (TF) activity differences analysis for plaque fibroblasts and T+B cell-enriched lymphoid aggregates following leuprolide treatment. Analyses comprised 33,026 and 3,830 spatial profiles within pathologist-defined coronary plaque and normal vascular regions, respectively, obtained from four murine hearts (two untreated and two leuprolide-treated). EMT; epithelial/endothelial-to-mesenchymal transition.

**Supplementary Figure 4.**
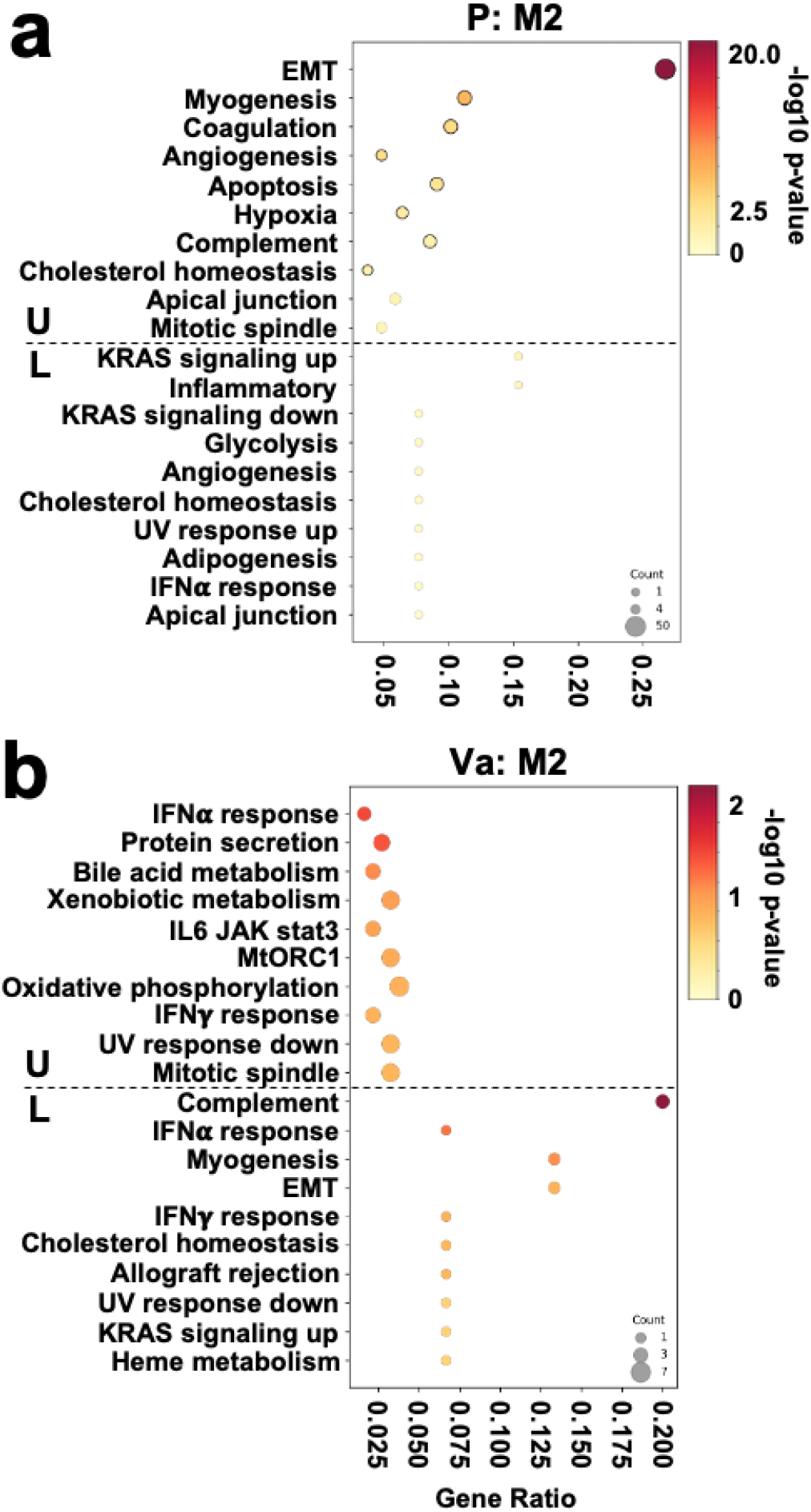
M2-like macrophages exhibit distinct remodeling programs in coronary plaque and normal vasculature. a,b, Hallmark pathway enrichment analysis of M2-like macrophages present in pathologist-defined coronary plaque (P; **a**) and histologically normal vasculature (Va; **b**) of untreated (U) and leuprolide-treated (L) murine hearts. Experimental design is shown in **Figure 1a**. Analyses comprised M2-like macrophage profiles derived from 33,026 and 3,830 spatially resolved cellular profiles within plaque and normal vascular regions, respectively, obtained from four murine hearts (two untreated and two leuprolide-treated).

**Supplementary Figure 5.**
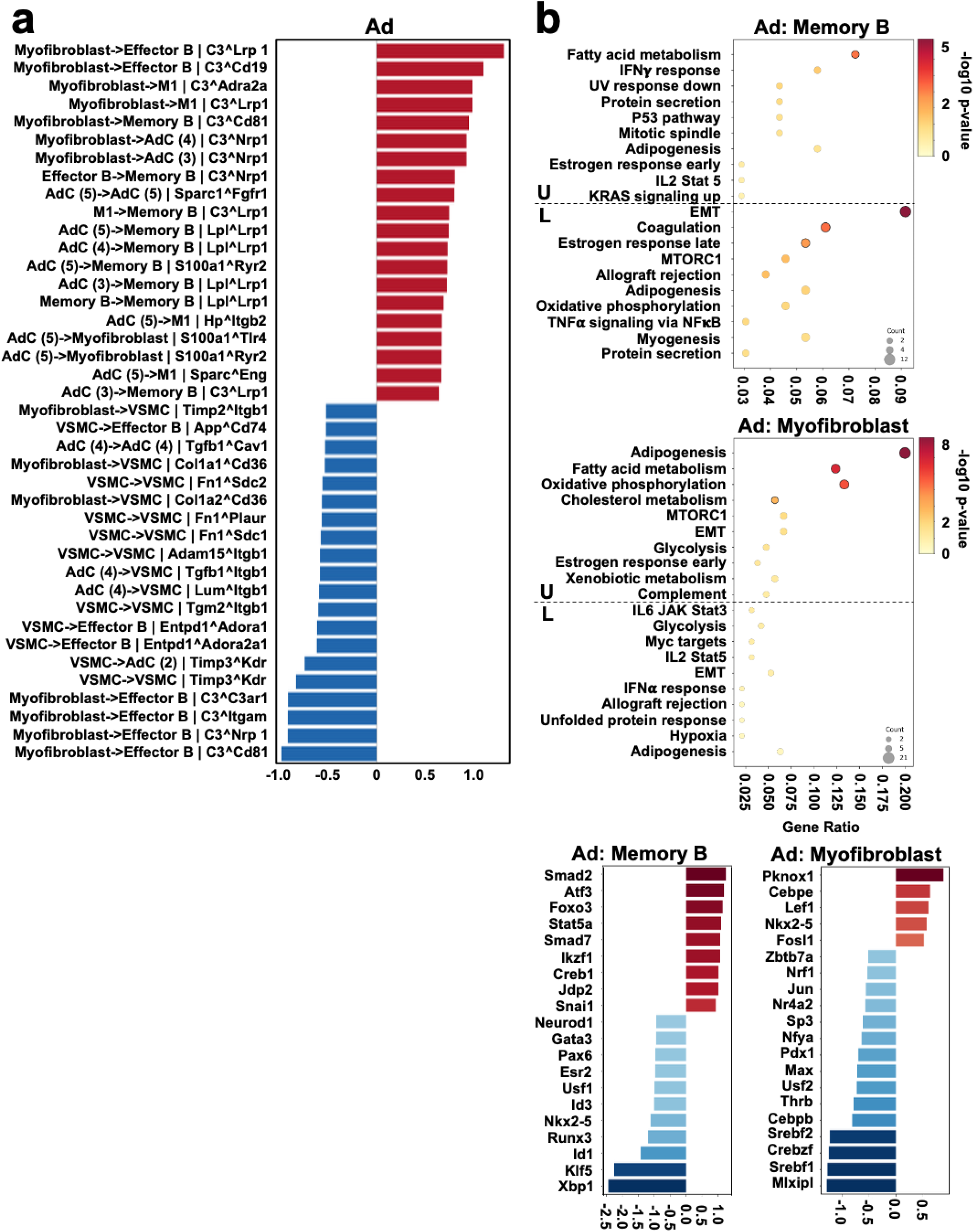
Leuprolide remodels adipocyte-B cell-myofibroblast communication in cardiac adipose tissue. **a,** Differential LIANA-predicted ligand-receptor interactions among cellular populations within cardiac adipose tissue (Ad) following leuprolide treatment (L) relative to untreated controls (U). Experimental design is shown in **Figure 1a**. Interactions were inferred from spatial transcriptomic expression of cognate ligand-receptor pairs. Bars show the treated-minus-untreated difference in LIANA consensus magnitude for the top 20 leuprolide-high (red color) and top 20 untreated-high (blue color) sender-receiver ligand-receptor combinations. Arrows indicate the direction of predicted signaling from ligand-expressing sender populations to receptor-expressing receiver populations. **b,** Hallmark pathway enrichment analysis and CollecTRI/ULM transcription factor (TF) activity of memory B cells and myofibroblasts present in cardiac adipose tissue of untreated and leuprolide-treated hearts. TF bars show treated-minus-untreated differences in mean profile-level ULM activity for the 20 TF with the largest absolute differences. Analyses comprised cellular profiles derived from 12,229 spatial profiles within pathologist-defined cardiac adipose regions obtained from four murine hearts (two untreated and two leuprolide-treated).

**Supplementary Figure 6.**
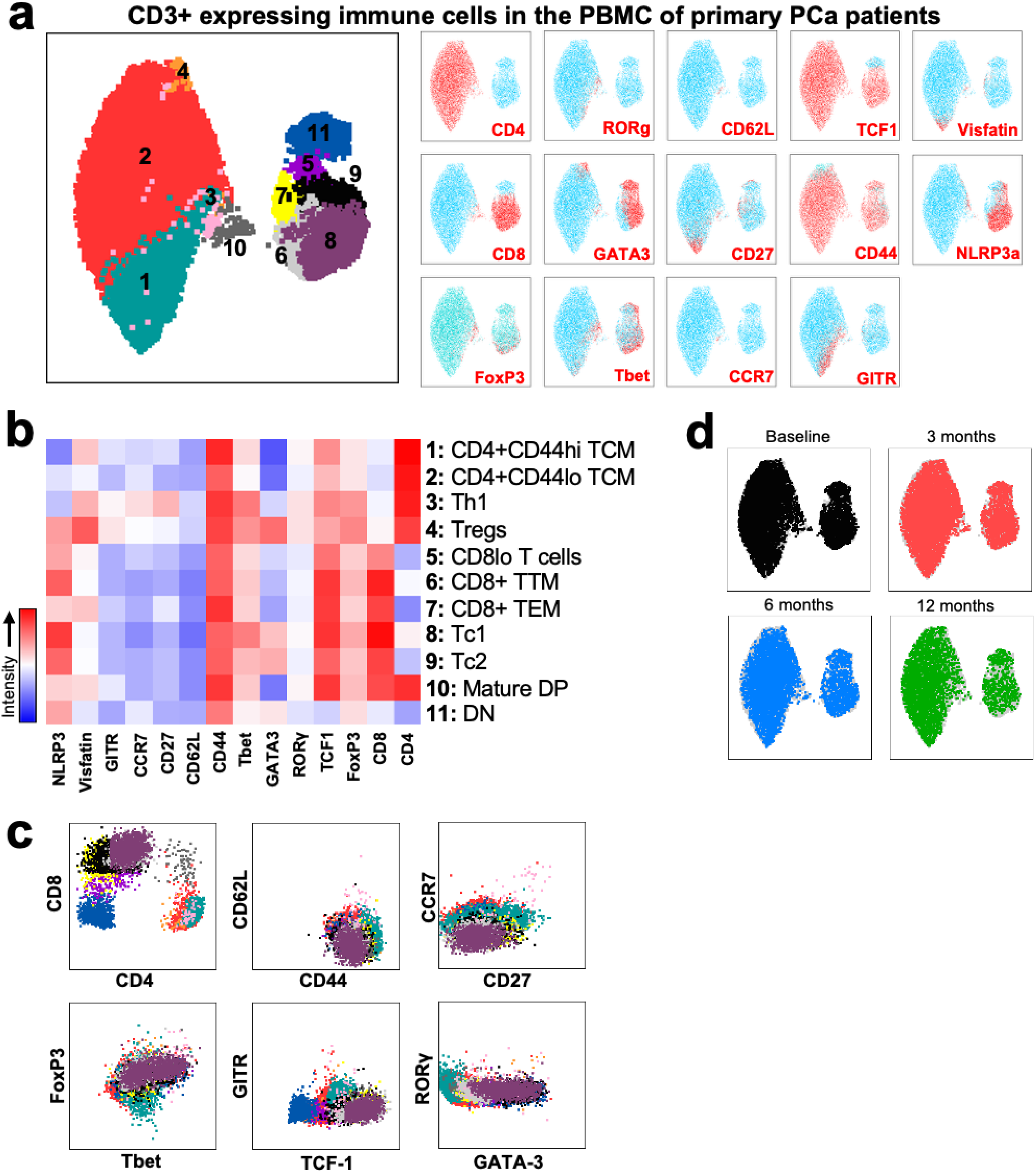
High-dimensional flow cytometry resolves 11 circulating T cell phenotypes in patients with primary prostate cancer. **a,** UMAP embeddings CD3**^+^**T cells of peripheral blood mononuclear cells (PBMC) collected from patients with primary prostate cancer enrolled in the CoPro study. Experimental design is shown in **Figure 4a**. Unbiased clustering based on the indicated 14 lymphoid markers resolved 11 distinct phenotypic T cell populations (left). Feature plots (right) show the relative expression of individual markers across the UMAP space used to characterize and annotate each population. **b,** Heatmap showing relative expression of the indicated phenotypic markers across the 11 T cell populations, supporting annotation of TCM; Central memory T cells, TTM; Transitional memory T cells, TEM; Effector memory T cells, DP; Double positive (CD4/CD8)**^+^**T cells, DN; Double negative (CD4/CD8)**^-^**T cells, Th1; Helper T cell type 1, Tregs; Regulatory T cells, Tc1; Cytotoxic T cell type 1 and Tc2; Cytotoxic T cell type 2. **c,** Flow-cytometry gating strategy illustrating marker combinations used to independently resolve and quantify the 11 T cell populations identified by high-dimensional clustering. **d,** UMAP visualization of circulating T cell populations according to longitudinal sampling time point at baseline and after 3 months, 6 months and 12 months of leuprolide treatment. Analyses included PBMC specimens from 25 patients with primary prostate cancer enrolled in the CoPro study across available longitudinal sampling time points.

**Supplementary Figure 7.**
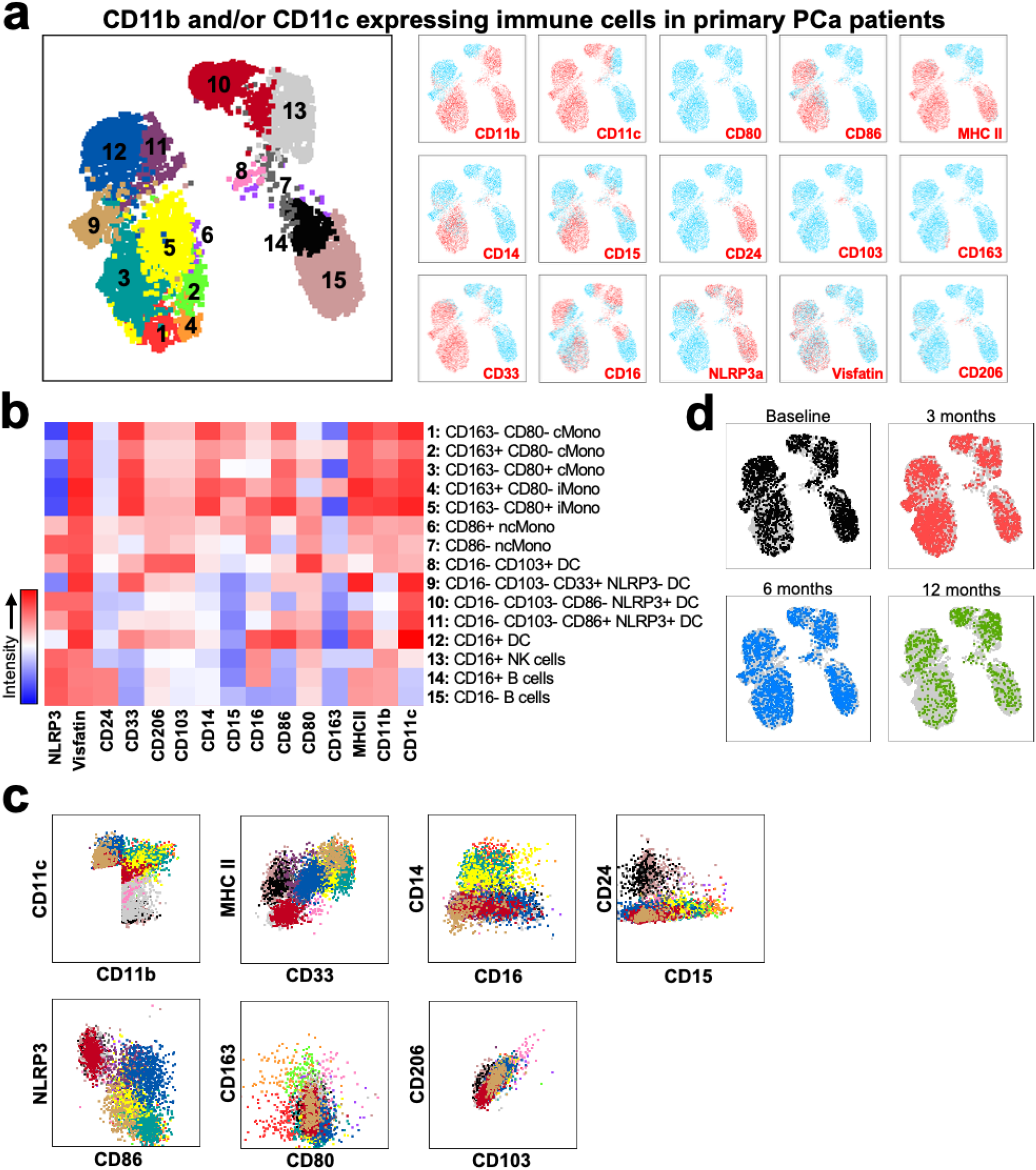
High-dimensional flow cytometry resolves circulating myeloid phenotypes in patients with primary prostate cancer. **a,** UMAP embedding CD11b**^+^**and/or CD11c**^+^**circulating immune cells of peripheral blood mononuclear cells (PBMC) collected from patients with primary prostate cancer enrolled in the CoPro study. Experimental design is shown in **Figure 4a**. Unbiased clustering based on the indicated 15 phenotypic markers resolved 15 distinct immune-cell populations (left). Feature plots (right) show the relative expression of individual markers across UMAP space used to characterize and annotate each population. **b,** Heatmap showing relative expression of the indicated markers across the 15 immune cell populations, supporting annotation of classical (cMono), intermediate (iMono) and non-classical (ncMono) monocyte states, dendritic cell (DC) populations, natural killer (NK) cells and B cells. **c,** Two-marker phenotypic classification strategy illustrating the marker combinations used to distinguish and quantify the 15 immune cell populations identified by high-dimensional clustering. **d,** UMAP visualization of the CD11b/CD11c-defined immune populations according to longitudinal sampling time point at baseline and after 3 months, 6 months and 12 months of leuprolide treatment, illustrating the distribution of major myeloid and innate immune phenotypes during therapy. Analyses included PBMC specimens from 25 patients with primary prostate cancer enrolled in the CoPro study across available longitudinal sampling time points.

**Supplementary Figure 8.**
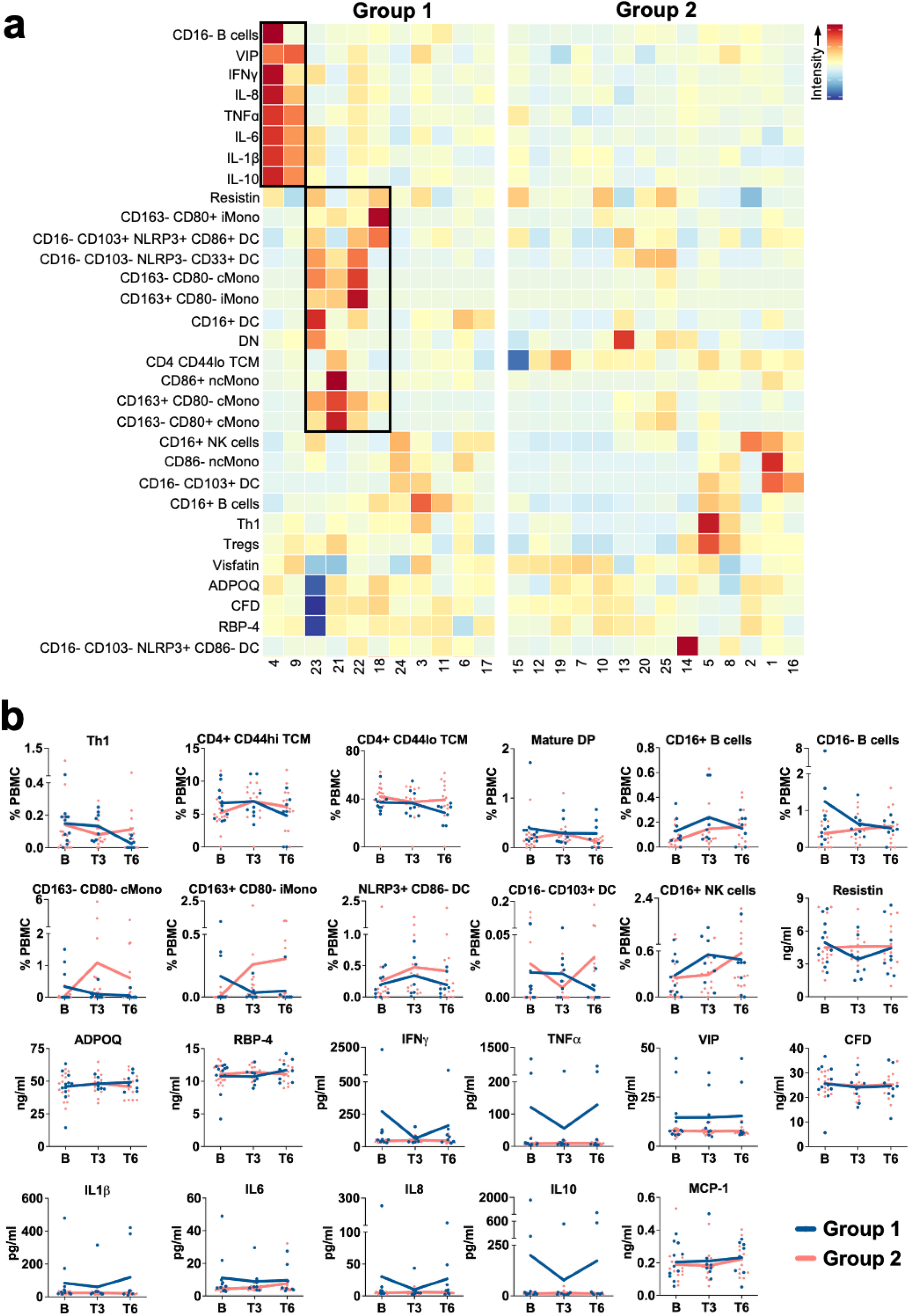
Myeloid and adipocytokine profiles do not distinguish the circulating T cell-defined patient phenotypes. **a,** Baseline circulating immune cell frequencies and adipocytokine concentrations in patients with primary prostate cancer stratified into Group 1 and Group 2 according to the unsupervised clustering analysis shown in **Figure 4b**. Individual patient values are shown for the indicated lymphoid, myeloid and soluble parameters. **b,** Longitudinal changes in the indicated circulating lymphoid and myeloid cell populations and adipocytokine concentrations in immune-defined Group 1 and Group 2 patients at baseline (B) and after 3 months (T3) and 6 months (T6) of leuprolide treatment. These analyses complement the populations showing significant treatment-associated changes in **Figure 4c,d** and demonstrate that the remaining circulating immune and soluble parameters exhibit limited group-specific or longitudinal alterations. Analyses included up to 25 patients with primary prostate cancer enrolled in the CoPro study; the number of evaluable patients at each longitudinal time point is indicated in the corresponding panels. Individual patient values are shown in **a**, and quantitative longitudinal data in **b** are presented as indicated. Data represents as mean±s.e.m. Statistical analyses for longitudinal immune and adipocytokine measurements are done using student T-test for panel **b**.

**Supplementary Figure 9.**
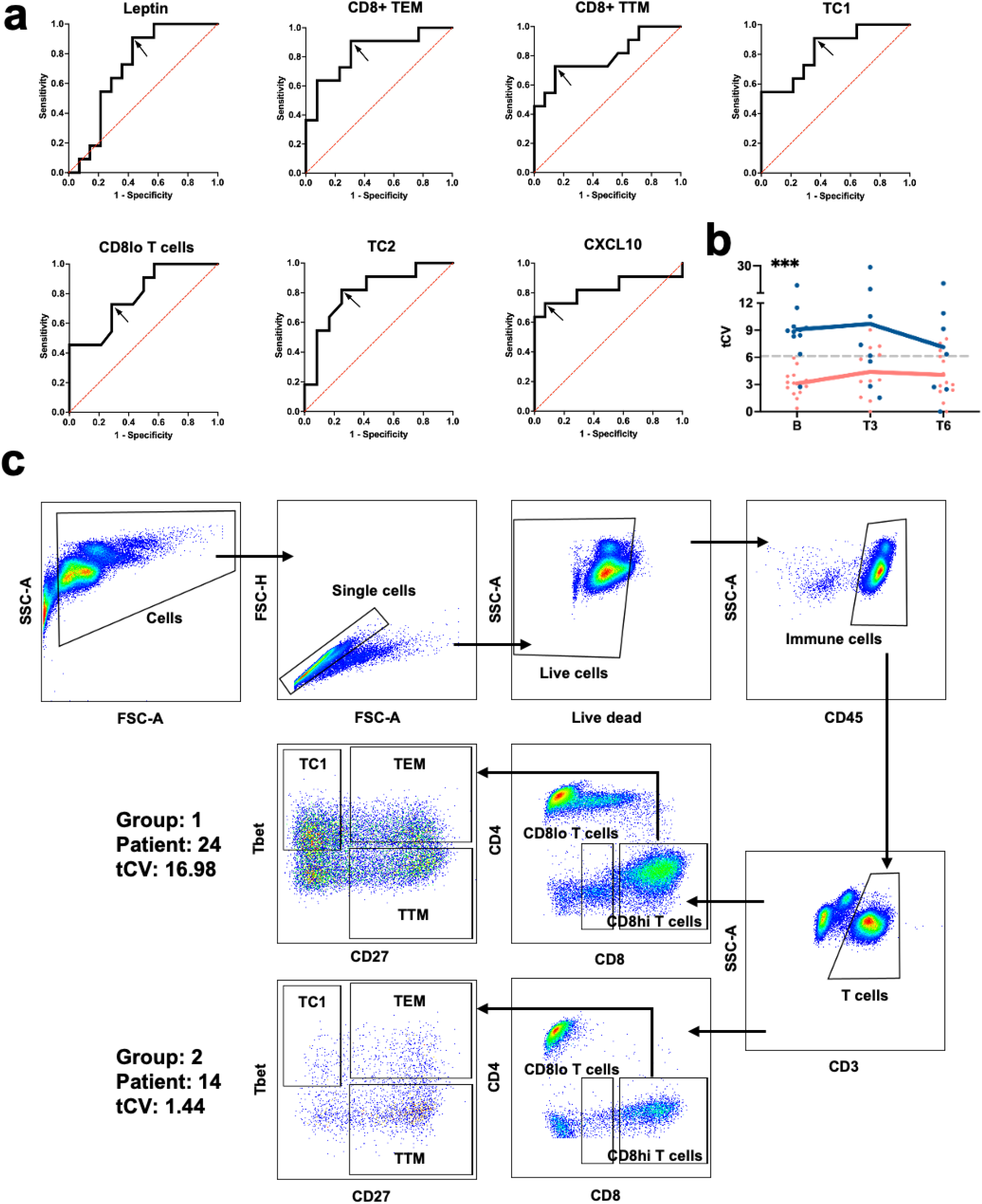
CD8^+^T cell components define and longitudinally maintain the tCV-associated immune phenotype. **a,** Receiver operating characteristic (ROC) analyses evaluating the individual contributions of the indicated circulating immune cell populations to discrimination of the baseline immune-defined patient groups identified in **Figure 4b**. Individual CD8**^+^**T cell populations and adipocytokines were evaluated using univariable logistic regression and compared with the composite multivariable tCV signature described in **Figure 4f**. Optimal thresholds were determined using the Youden index. **b,** Longitudinal distribution of the composite tCV score at baseline (B) and after 3 months (T3) and 6 months (T6) of leuprolide treatment in patients stratified according to the baseline immune-defined groups. The analysis evaluates the stability of the CD8**^+^**T cell-associated signature during treatment rather than *de novo* CVE prediction at individual time points. **c,** Representative flow-cytometry plots show the CD8**^+^**T cell subsets contributing to the composite signature in two patients, and selected to illustrate high and low baseline tCV scores. The contributing populations include Tc1 (cytotoxic type 1 CD8**^+^**T cells), CD8**^+^**effector-memory (TEM), CD8**^+^**transitional-memory (TTM) and CD8^lo^T cell states. Analyses included up to 25 patients with primary prostate cancer enrolled in the CoPro study, with the number of evaluable samples at each longitudinal time point indicated in the corresponding panels or source data. Data represents as mean±s.e.m. Statistical analyses were done using student T-test for panel **b** and presented; ***p<0.001.

**Supplementary Table 1.** Cardiovascular events and immune-defined phenotypes in patients with primary prostate cancer enrolled during CoPro study.

| *Patient # | Age | Pre-leuprolide CVE and risk factor | On-leuprolide CVE | Type of CVE | Group |
| --- | --- | --- | --- | --- | --- |
| 2 | 77 | Sinus node dysfunction | None | None | 2 |
| 4 | 69 | Tachycardia with elevated coronary artery calcification with smoking history | None | None | 2 |
| 7 | 65 | Paroxysmal atrial fibrillation and enlarged thoracic aorta with Percutaneous coronary intervention | None | None | 2 |
| 10 | 65 | Elevated coronary artery calcification with smoking history | Unspecified angina with native coronary artery atherosclerosis | M-CVE | 2 |
| 12 | 62 | Cardiomegaly, diastolic heart failure, aortic valve disorder with smoking history | Angina | M-CVE | 2 |
| 14 | 67 | Carotid artery stenosis | Sub-clinical coronary atherosclerosis, hypertension | M-CVE | 2 |
| 15 | 64 | Right bundle branch block with severe calcified plaque volume | None | None | 2 |
| 16 | 72 | Moderate calcified plaque in the left anterior descending coronary artery | None | None | 2 |
| 18 | 79 | Rheumatic disorder of mitral and aortic valves, cardiomyopathy and thoracic aorta ectasia with defibrillator installed and had smoking history | None | None | 2 |
| 19 | 79 | Mitral regurgitation, ischemic cardiomyopathy, chronic systolic heart failure with coronary artery bypass grafting and had smoking history | None | None | 1 |
| 20 | 70 | Atrial fibrillation | None | None | 2 |
| 22 | 61 | None | Premature ventricular complexes and nonspecific ST abnormality | T-CVE | 1 |
| 24 | 75 | None | Nonischemic cardiomyopathy with reduced left ventricle ejection fraction | T-CVE | 1 |
| 25 | 76 | None | Angina with negative cardiac work-up indicating reaction to COVID booster shot | M-CVE | 1 |
| **M-CVE** = Multifactorial cardiovascular events (CVE); physician can expect or predict the CVE based on patient history of cardiovascular diseases or risk factors<br>**T-CVE** = Treatment-associated cardiovascular events; Unexpected by physician based on history of cardiovascular diseases or risk factors<br>*CoPro trial enrolled 25 patients and retrospectively able to collected CVE from 14 patients. |  |  |  |  |  |

